# Multi-Scale Kinetics Modeling and Advanced Assay for mRNA-Lipid Nanoparticle Potency Assessment

**DOI:** 10.1101/2025.09.29.679406

**Authors:** Yuling Yang, Yuchen Qiu, Keqi Wang, Yifang Liu, Gautam Sanyal, Paul C. Whitford, Sara H. Rouhanifard, Wei Xie

## Abstract

mRNA lipid nanoparticle (mRNA-LNP) technology has attracted global attention, especially in vaccine development, due to its superior delivery efficiency, molecular stability, and safety profile. However, evaluating mRNA-LNP potency—defined as the quantifiable biological response elicited by the product—remains costly and time-consuming when relying solely on *in vitro* experiments. Rapid and reliable potency assessment is hindered by limited mechanistic understanding of delivery processes and sparse experimental data. To address these challenges, we present a mechanism-informed, multi-scale kinetic modeling framework that quantitatively captures the coupled dynamics across particle-level, cellular, and macroscopic scales. This model incorporates variability in LNP-cell interactions and accounts for critical factors such as dosage, LNP and cell size distributions, and cell proliferation dynamics—all of which influence delivery efficiency and response variability. By integrating advanced multi-omics assays—such as single-molecule fluorescent in situ hybridization (smFISH), which enables single-cell resolution of mRNA and protein expression—our framework leverages heterogeneous, multi-scale data to support mechanistically grounded modeling of mRNA delivery and enable robust predictions of therapeutic potency.

**Graphical Abstract:** 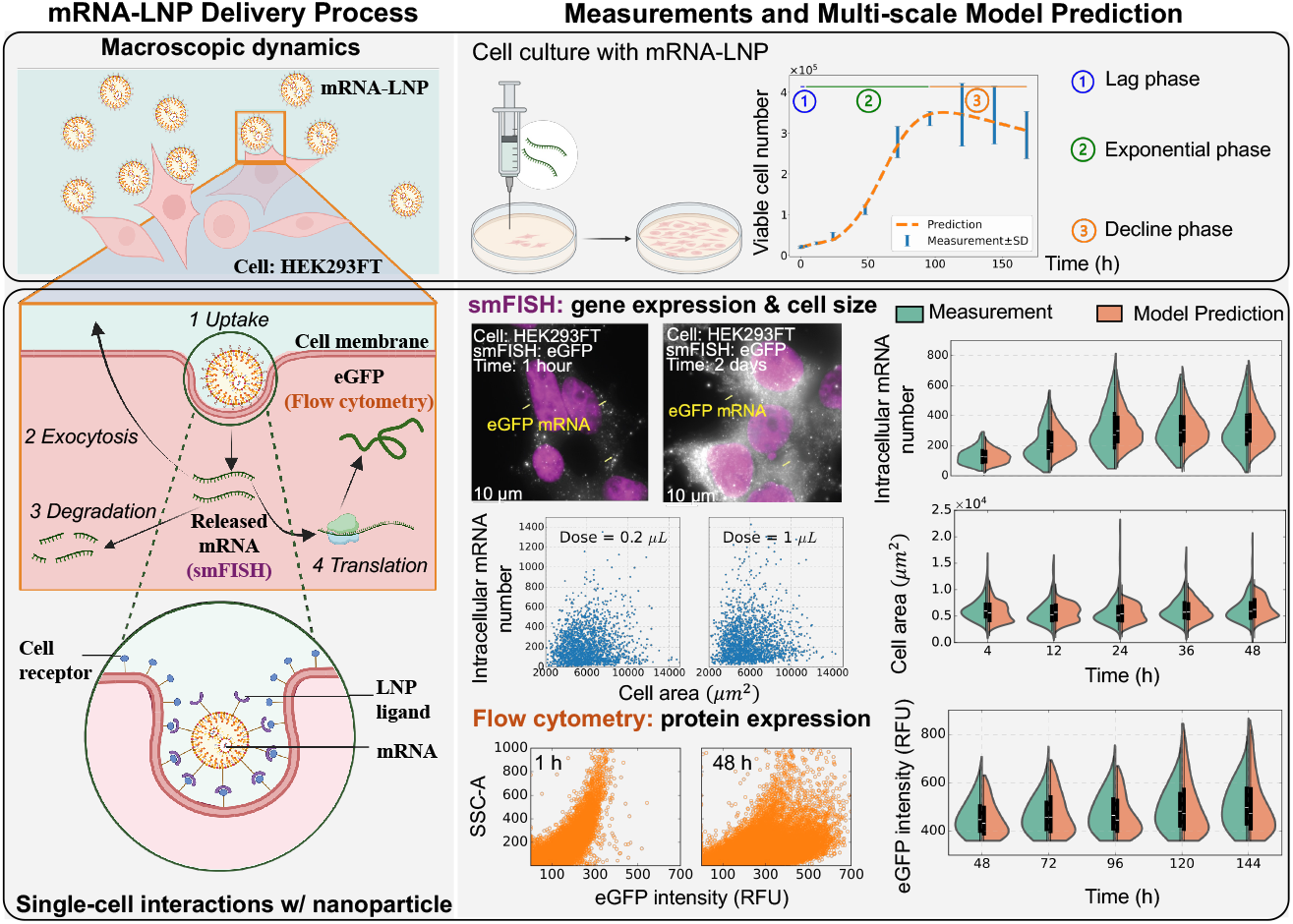

## Introduction

Due to the critical role in the development of COVID-19 vaccines, the lipid nanoparticle (LNP) mediated mRNA delivery system has attracted global attention. Designed with optimal size and specific surface properties, LNPs are beneficial for stability and delivery efficiency for nucleic acid therapies [1, 2]. Despite significant progress in mRNA–LNP technology, the development of rapid and reliable potency assessment tools remains hindered by an incomplete understanding of the complex and variable mechanisms that regulate RNA delivery and expression.

Potency refers to the measurable biological response elicited by a drug product and is critically dependent on its functional integrity [3]. This study focuses on cell-based potency assessment of mRNA–LNP systems by quantifying transfection efficiency and evaluating the expression levels of the target antigen protein. From a kinetic perspective, ligand-receptor-mediated endocytosis plays a central role in determining the transfection rate, as ligands on the LNP surface initiate receptor-mediated endocytosis and subsequent internalization by the target cell [4]. The dynamics of the delivery process is influenced by several factors, including dose, LNP size, mRNA-LNP concentration that have not been uptake yet, and the densities of ligands on nanoparticles and cell surface receptors [5]. Additionally, stochastic variables such as cell surface area and viability, as well as the size and surface charge of LNPs, contribute to cell-to-cell variability in mRNA–LNP uptake [6, 7, 8]. As a result, gene and protein expression exhibit substantial heterogeneity across cells, driven not only by differences in LNP uptake but also by stochastic influences such as mRNA degradation, intracellular release kinetics, and innate immune activation [9]. Therefore, cell-to-cell heterogeneity can influence the overall response to mRNA-LNP delivery by causing variable transfection efficiency and protein expression across different cells, which may lead to inconsistent immune activation and therapeutic outcomes in the human body [10, 11].

A challenge arises from studying the interactive effect from multiple sources of variations on the potency. Specifically, how do we quantify the contributions from various sources of uncertainty on cell-based potency? This information can guide quality control of mRNA-LNP systems and improve the consistency of potency. Although recent studies have begun to model the variability in cell size [7], stochastic fluctuation of particle uptake [12], and gene expression [13], a comprehensive investigation into how these factors influence critical quality attributes (CQAs), such as the effect of LNP size on delivery efficiency and cell-based potency, remains lacking. Such an analysis is essential to inform the optimal design of mRNA-LNP products and to minimize the reliance on extensive physical experimentation.

LNP size influences uptake in two ways: LNP endocytosis time and mRNA loading. The size of the LNP determines endocytosis rate by the balance between the adhesion energy and the deformation energy of the cell membrane [6, 14]. When the adhesion strength and ligand density are sufficiently high, receptor diffusion can become the dominant, rate-limiting step of endocytosis process. By developing a front-tracking diffusion model, the size-dependent endocytosis rate of nanoparticle (NP) is quantified and various scaling analyses are performed in existing studies [5, 6, 15, 16]. In addition, larger LNPs tend to carry more mRNA per particle [17, 18]. Thus, for a fixed total mRNA dose, increasing the size of the LNP reduces the concentration of the particles and alters LNP-cell interactions. Understanding these coupled effects is critical, as they collectively determine how LNP size influences delivery process dynamics and potency variability. However, to date, the interplay between endocytosis rate and mRNA concentration at cellular and macroscopic levels has not been systematically characterized.

Another major challenge in *in vitro* assessments is the limited understanding of mRNA-LNP delivery kinetics in proliferating cells. Commonly used cell lines for *in vitro* mRNA-LNP studies, such as HEK293 and its derivatives [19, 20, 21, 22], as well as HeLa cells [23, 24, 25], typically have doubling time ranging from 24 to 48 hours [23, 19, 26, 27]. Cell proliferation affects the dynamics of mRNA-LNP delivery through two primary mechanisms: it alters the LNP uptake capacity of individual cells and contributes to the dilution of intracellular molecules due to cell division. Since each cell has a finite number of surface receptors, the cell density influences how many LNPs can be taken by cells simultaneously. Under conditions of elevated cell density and constrained LNP supply, individual cells receive fewer nanoparticles as a result of resource depletion and intercellular competition. In terms of intracellular molecule dilution, endogenous molecules are typically replicated during each cell cycle and segregated into two daughter cells during cell division [28]. However, mRNA-LNP vaccines are considered safe due to the non-replicating and non-integrating nature of the delivered mRNA [29]. These properties result in progressive dilution of intracellular mRNA in proliferating cells, which can contribute to variability in delivery efficiency and therapeutic potency over time. To the best of our knowledge, there is a lack of specific studies focusing on modeling mRNA-LNP delivery in proliferating cells.

The final challenge lies in performing single-cell mRNA measurements. For mRNA-LNP systems, *in vitro* potency assays are typically used to assess transfection efficiency and the generation of the target (antigen) protein [23], e.g., flow cytometry [13, 30, 22, 31] and quantitative reverse transcription polymerase chain reaction (qRT-PCR) [32, 33, 34]. Conventional approaches, such as bulk assays like qRT-PCR, lack the resolution to capture mRNA dynamics at the single-cell level. Although live-cell imaging has been widely used to monitor single-cell expression of fluorescent proteins [12, 35, 36, 37], therapeutic mRNAs encapsulated in LNPs typically do not include fluorescent reporter genes. This is because the therapeutic goal is to deliver a minimal cargo—namely, the mRNA itself—without additional genetic elements, making direct tracking of protein expression infeasible.

To address these challenges, in this study, we collected time-course measurements on mRNA and protein expression, aiming for better scalability and industrial relevance. In particular, three types of assays, including single-molecule fluorescent in situ hybridization (smFISH), flow cytometry, and qRT-PCR, provide time-course measurements on the distribution changes of single-cell mRNA and protein expression, as well as mRNA concentration in the extracellular solution prior to cellular uptake. By integrating heterogeneous experimental data collected across multiple scales, we developed a multi-scale stochastic kinetic model that characterizes single-cell dynamics and the variability in interactions between LNPs and cells. This model enables mechanistic insights into RNA delivery and expression processes at molecular, cellular, and macroscopic levels. While previous studies have employed deterministic approaches to model individual subprocesses within the delivery system [38, 39, 40, 41], they often fail to account for cell-to-cell variability in potency. Our model overcomes this limitation by integrating intrinsic stochasticity into single-cell dynamics, enabling a more faithful representation of biological heterogeneity.

The objective of the proposed framework is to improve the prediction of cell-based potency by leveraging multi-scale kinetic modeling and mechanism learning. Ligand-receptor interactions are modeled using a Monod kinetic formulation, where binding rates are influenced by factors such as extracellular mRNA-LNP concentration, LNP size, cell surface area and growth rate. Upon binding, mRNA-LNP complexes are gradually internalized by cells. The impact of LNP size and mRNA loading on the internalization rate is captured through a physics-informed, front-tracking diffusion model. Upon internalization, mRNA-LNPs follow one of two pathways: exocytosis or intracellular mRNA release. Exocytosis occurs when the intracellular concentration of mRNA rises and reaches a saturation threshold. Alternatively, released mRNA is translated into the target protein at a rate proportional to its intracellular copy number. The model also incorporates dilution effects due to cell proliferation and division, ensuring an accurate representation of dynamic cellular processes and changes in mRNA and protein expression over time.

The proposed multi-scale kinetic model effectively captures cell-to-cell variability in both delivery and expression processes. It accounts for diverse sources of heterogeneity stemming from LNP and cellular properties, such as LNP size and cell surface area. Experimental validation confirms that the model can reliably predict cell-based potency with minimal input, offering a rapid and accurate assessment tool. As a result, this framework not only supports quality assurance efforts but also provides mechanistic insights to guide the optimization of mRNA-LNP delivery strategies across heterogeneous target cell populations.

## Materials and Methods

In this section, we first introduce the multi-scale kinetic model of mRNA-LNP delivery and expression, and then elaborate the multi-omics data collection and analysis.

### Mechanism-Informed Multi-Scale Kinetic Model

To characterize the mechanisms of mRNA-LNP delivery and expression across molecular, cellular, and macroscopic scales, we propose a Mechanism-Informed Multi-Scale (MIMS) model with a hierarchical and modular architecture. In the following sections, we detail the core components of MIMS, including its structural design, underlying assumptions, dynamic modeling of regulatory mechanisms in subprocesses, and model inference approach.

### Hierarchical and Modular Design of MIMS

The proposed multi-scale kinetic model aims to elucidate the underlying regulatory mechanisms governing mRNA-LNP delivery and expression across nanoparticle, cellular, and macroscopic scales; see a cartoon illustration in Figure 1. The rate that cells metabolize mRNA-LNPs is influenced by the extracellular environment, including nutrient availability and the accumulation of metabolic waste, as well as by cell growth, division, and cell death. Consequently, the regulatory mechanisms governing mRNA-LNP delivery and expression—captured by the parameters of the kinetic model denoted by ***θ***—are dependent on the cell’s metabolic phase, such as the lag, exponential growth, and decline phases.

**Fig. 1:**
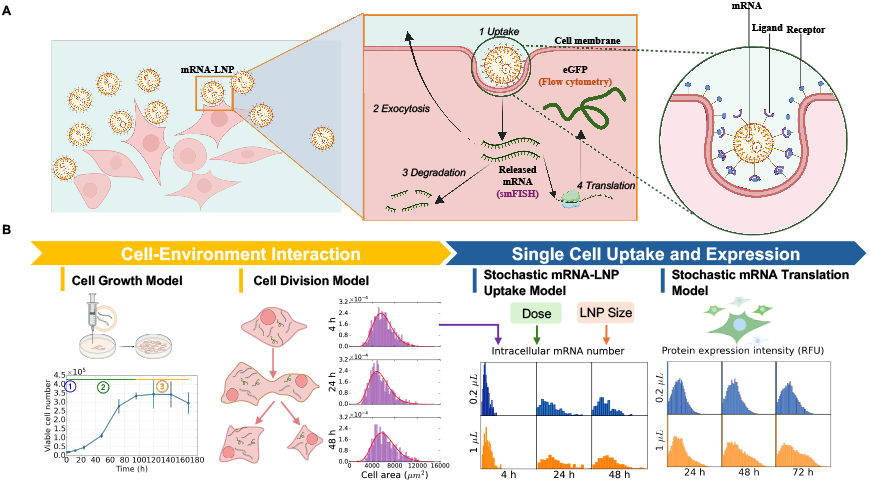
Hierarchical and modular architecture of the proposed Mechanism-Informed Multi-Scale model (MIMS). (A) A schematic illustration of the mRNA-LNP delivery and expression process, highlighting key events: (1) cellular uptake via ligand–receptor binding, (2) mRNA exocytosis, (3) mRNA degradation, and (4) translation. (B) The cell growth module captures cell density dynamics across three phases: ➀ lag phase, ➁ exponential growth phase, and ➂ decline phase. The cell division module models stochastic, cell area–dependent division behavior. The uptake module estimates the dynamics of intracellular mRNA copy number based on extracellular mRNA concentration, LNP content, and the sizes of both the cell and the nanoparticle. In contrast, the translation module predicts protein expression levels from intracellular mRNA abundance.

The proposed multi-scale kinetic model has a hierarchical structure across nanoparticle, cellular, and macroscopic scales. We summarize the variables of interest in Table 1. The cell culture duration is discretized into *H* uniform time intervals of length Δ*t*, with discrete time points denoted as *t*_*h*_ = *h*Δ*t* for *h* = 0, 1, …, *H*. To estimate the expected loading efficiency of LNPs, we quantified the initial extracellular mRNA concentration using qRT-PCR. The system inputs at time zero included the dose, the measured extracellular mRNA concentration *s*_mRNA.*ex*_(0) (molecules), and the initial size distribution of the LNPs *s*_LNP.size_(0) (nm).

**Table 1.** Key variables in the MIMS framework.

| Scale | Symbol | Type and Composition | Description |
| --- | --- | --- | --- |
| Nanoparticle | $\mathbf{s}_{\text{LNP}}(t_h)$ | Vector:<br>$[s_{\text{LNP.size}}(t_h), s_{\text{LNP.load}}(t_h)]$ | Nanoparticle size $s_{\text{LNP.size}}(t_h)$ and mRNA loading $s_{\text{LNP.load}}(t_h)$. |
| Single Cell | $\mathbf{s}_{\text{Molecule}}(t_h)$ | Vector:<br>$[s_{\text{mRNA.in}}(t_h), s_{\text{Protein}}(t_h)]^\top$ | Intracellular molecular states: mRNA copy number $s_{\text{mRNA.in}}(t_h)$ and protein expression intensity $s_{\text{Protein}}(t_h)$. |
| | $\mathbf{s}_{\text{Cell}}(t_h)$ | Vector:<br>$[A(t_h), z(t_h), \mathbf{s}_{\text{Molecule}}(t_h)^\top]^\top$ | Cell state: cell area $A(t_h)$, cell cycle phase $z(t_h)$, and molecular contents $\mathbf{s}_{\text{Molecule}}(t_h)$. |
| | $\mathbf{v}(t_h)$ | Vector | Single-cell reaction rates associated with delivery and expression subprocesses at time point $t_h$. |
| Macroscopic | $X(t_h)$ | Scalar | Viable cell density at time $t_h$. |
| | $s_{\text{mRNA.ex}}(t_h)$ | Scalar | Concentration of extracellular mRNAs encapsulated in LNPs. |

At the nanoparticle scale, the state of each LNP, represented ***s***_LNP_(*t*_*h*_) = [*s*_LNP.size_(*t*_*h*_), *s*_LNP.load_(*t*_*h*_)], is specified by particle size *s*_LNP.size_(*t*_*h*_) (nm) and the mRNA loading *s*_LNP.load_(*t*_*h*_). At the single-cell scale, we track the intracellular molecular state, i.e., ***s***_Molecule_(*t*_*h*_) = [*s*_mRNA.*in*_(*t*_*h*_), *s*_Protein_(*t*_*h*_)]^*⊤*^, consisting of mRNA copy number *s*_mRNA.*in*_(*t*_*h*_) (molecules per cell) and protein expression intensity *s*_Protein_(*t*_*h*_) in terms of Relative Fluorescence Unit (RFU) per cell. To capture single-cell states in addition to gene and protein expression, we incorporate cell surface area *A*(*t*_*h*_) and cell cycle phase *z*(*t*_*h*_) (i.e., G1, S, G2, and M), both of which influence cell division behavior. Thus, we concatenate *A*(*t*_*h*_), *z*(*t*_*h*_), and ***s***_Molecule_(*t*_*h*_) into a single-cell state vector, i.e., ***s***_Cell_(*t*_*h*_) = [*A*(*t*_*h*_), *z*(*t*_*h*_), ***s***_Molecule_(*t*_*h*_)^*⊤*^]^*⊤*^. At the macroscopic scale, the environmental conditions, including the viable cell density denoted as *X*(*t*_*h*_) (cells) and the extracellular mRNA concentration denoted as *s*_mRNA.*ex*_(*t*_*h*_) (molecules), influence cell growth and activity to process mRNA-LNPs. Here, we focus on the extracellular mRNAs encapsulated in LNPs, as naked RNAs are unable to enter cells due to electrostatic repulsion between their negatively charged phosphate backbone and the negatively charged cell membrane. Therefore, at any time *t*_*h*_, the multi-scale state variables are represented as,

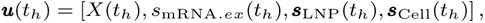

with the distributions of ***s***_LNP_(*t*_*h*_) and ***s***_Cell_(*t*_*h*_) characterizing the variability in the attributes of individual lipid nanoparticles and cells, respectively.

The modular design of the proposed MIMS model comprises single-cell kinetic modules that represent cell growth and the subprocesses involved in mRNA-LNP delivery and expression. Cell growth dynamics influence both cell area-dependent division and the intracellular distribution of molecules between daughter cells. Specifically, cell division events are simulated using a Markov chain model where the transition probabilities are functions of both cell area *A*(*t*_*h*_) and cell cycle phase *z*(*t*_*h*_). At the macroscopic scale, cell division events are aggregated to model the dynamics of cell population density *X*(*t*_*h*_), i.e.,

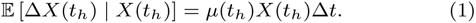

The cell density growth rate *µ*(*t*_*h*_) serves as a proxy for cell viability and is modulated by extracellular conditions such as nutrient availability and the accumulation of metabolic waste. This time-dependent growth rate is used to infer the kinetic model parameters ***θ***. Specifically, *µ*(*t*_*h*_) enables identification of distinct cell culture phases—such as lag, exponential growth, and decline phases (see Figure 1)—allowing the model to assign phase-specific parameter values that capture dynamic changes in cellular behavior across these phases.

The single-cell delivery and expression process is modularized into *c* = 4 distinct subprocesses: mRNA-LNP uptake, mRNA exocytosis, translation, and degradation. The reaction rates associated with these subprocesses are represented by the vector ***v*** = [*v*_up_, *v*_exo_, *v*_TL_, *v*_deg_]^*⊤*^, and are functions of the system state at time *t*_*h*_, denoted by ***u***(*t*_*h*_). Their influence on the state dynamics *{****u***_*t*_*}* is captured using a modular framework:

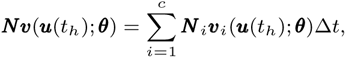

where ***N*** is an *m×n* stoichiometry matrix defining the structure of the delivery and expression reaction network, with *m* denoting the number of molecular species and *n* the number of reactions. The (*p, q*)-th element of ***N***, denoted as ***N*** (*p, q*), specifies the number of molecules of the *p*-th species consumed (negative value) or produced (positive value) in each occurrence of the *q*-th reaction.

These values serve as the stoichiometric coefficients for the *p*-th species in the rate equation of the *q*-th reaction.

#### Uptake

The single-cell delivery process begins with mRNA-LNP uptake, during which cell surface receptors bind to ligands on the LNPs. The internalization rate of each nanoparticle is size-dependent. During endocytosis, the number of ligand-receptor binding sites gradually increases, and the nanoparticle is internalized once all ligands are bound, as described in [5]. This uptake subprocess can be represented by the following reaction:

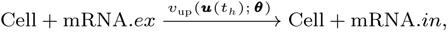

where mRNA.*ex* and mRNA.*in* represent extracellular and intracellular mRNAs. The mRNA uptake rate, denoted as *v*_up_(***u***(*t*_*h*_); ***θ***), is modeled as a function of the multi-scale system state ***u***(*t*_*h*_) and kinetic parameter set ***θ***.

#### Exocytosis

As cells internalize LNPs, they eventually reach a saturation state, at which point the released mRNAs are transported back into the extracellular environment via exocytosis. This exocytosis subprocess can be represented as follows:

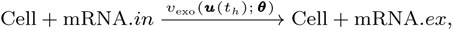

where *v*_exo_(***u***(*t*_*h*_); ***θ***) represents the exocytosis rate.

#### Translation

While some internalized mRNAs are exported back to the extracellular environment via exocytosis, others remain in the cytoplasm and serve as templates for synthesizing amino acid chains (denoted as AA) that form the target protein within ribosomes—a process known as translation [42]. The translation subprocess can be represented as follows:

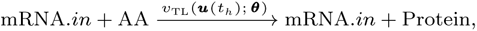

where *v*_TL_(***u***(*t*_*h*_); ***θ***) represents the translation rate.

#### mRNA Degradation

Some mRNAs eventually undergo degradation and are never utilized for translation, as represented below:

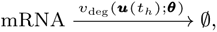

where *v*_deg_(***u***(*t*_*h*_); ***θ***) represents the mRNA degradation rate.

To model the evolution of intracellular molecular states for each cell, denoted by ***s***_Molecule_(*t*_*h*_), we employ a stochastic framework based on state transition probabilities. Specifically, the dynamics are governed by the following stochastic differential equation:

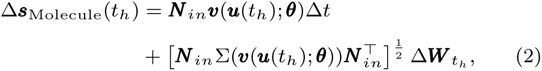

where ***N*** _*in*_ is a submatrix of the stoichiometry matrix ***N***, representing the reaction structure associated with intracellular molecules, Σ(***v***(***u***(*t*_*h*_); ***θ***)) is a diagonal covariance matrix capturing the intrinsic variability of each reaction, and Δ***W*** _*t*_*h* is a Gaussian random vector with mean zero and covariance matrix diag*{*Δ*t}* modeling thermodynamic fluctuations via Brownian motion. Eq. (2) enables the modeling of stochastic single-cell dynamics by accounting for biological variability arising from multiple factors, including cell proliferation and division, changes in cell area, ligand–receptor interactions, mRNA degradation, exocytosis, and translation. These components are described in detail in the following subsections.

In the following sections, we detail the modeling of potency dynamics and variability, along with the regulatory mechanisms governing reaction rates ***v***(*t*_*h*_). At the macroscopic scale, the rate of decrease in extracellular mRNA concentration is positively correlated with the mRNA-LNP uptake rate, up to a saturation threshold [23, 41]. Additionally, cell growth rate influences both the ligand-receptor binding kinetics and the dilution of intracellular molecules. At the single-cell scale, features such as cell area and membrane tension modulate endocytosis rates. At the particle scale, LNP-specific properties—including size, shape, and surface ligand density—govern the kinetics of endocytic uptake. The interplay across these hierarchical scales collectively determines the overall efficiency of cell-based potency.

### Model Assumptions

The following assumptions have been made for our model: (1) **Localization of mRNA-LNP delivery**. All delivery-related processes—including LNP uptake, mRNA release, and mRNA translation—are assumed to occur within the cytosol. This assumption excludes nuclear interactions associated with alternative pathways, thereby ensuring that endogenous cellular genes remain unaffected by exogenous mRNA. **(2) Immediate mRNA release upon internalization**. Once fully internalized by the cell, LNPs are assumed to release their mRNA loading immediately. This assumption is supported by the design of the lipid envelope, which promotes efficient intracellular mRNA release from both the LNP and the endosome—more effectively than cationic liposomes or polymers [1]. By assuming immediate release, the model simplifies the representation of intracellular kinetics by avoiding the complexity associated with delayed or partial release mechanisms. **(3) Negligible cell area growth during the G2-M phase**. Cell growth primarily occurs during the G1–S phase, while the G2–M phase is dedicated to preparing cellular components for division rather than increasing cell size [43, 44, 45]. Based on this biological insight, the model assumes negligible cell area growth during the G2–M phase, thereby simplifying the representation of cell division dynamics by treating cell area as static during this stage. **(4) mRNA loading of LNPs is proportional to LNP volume**. This assumption is supported by prior studies [18, 19], enabling indirect estimation of mRNA loading per nanoparticle. By quantifying the initial extracellular mRNA concentration using qRT-PCR and measuring the LNP size distribution via dynamic light scattering (DLS), the expected mRNA copy number per nanoparticle can be inferred based on its volume.

### Cell Growth Model

In the batch cell culture process, cell growth curves typically consist of multiple metabolic phases, including the lag phase, the exponential growth phase, and the decline phase [46]. A representative growth curve is shown in Figure 2C. This progression can be mathematically summarized as follows:

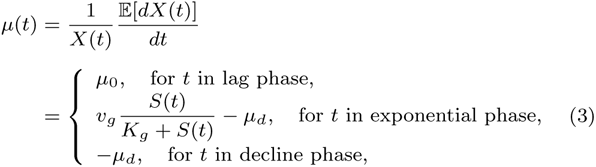

where *µ*(*t*) represents the time-dependent growth rate of cell density, *µ*_0_ represents cell growth rate in the lag phase, *µ*_*d*_ represents the constant cell death rate, *v*_*g*_ represents the max growth rate during exponential phase, *K*_*g*_ represents the Monod constant characterizing substrate-dependent growth, and *S*(*t*) is the concentration of a growth-related substrate in the extracellular environment at time *t*. Eq. (3) can be further discretized as Eq. (1)

**Fig. 2:**
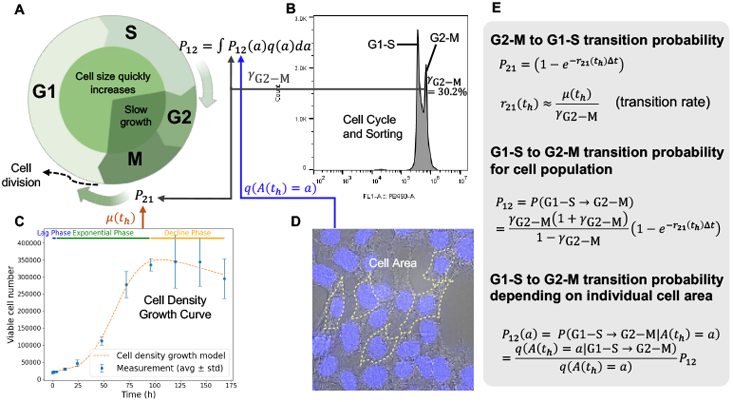
Integrative modeling cell cycle transitions and division dynamics as a function of cell area. (A) Schematic of the Markov chain model representing cell cycle transitions. The model divides the cell cycle into two principal phases—G1–S and G2–M—with transition probabilities *P*_12_ and *P*_21_ governed by the cell growth rate *µ*(*t*_*h*_), cell area *A*(*t*_*h*_), and the phase distribution parameter *γ*_G2-M_.(B) Flow cytometry analysis of cell cycle distribution using Vybrant™ DyeCycle™ Violet for DNA content staining reveals two distinct peaks, corresponding to populations of cells in the G1-S and G2-M phases. (C) The cell growth curve accurately captures the dynamics of cell proliferation and delineates distinct metabolic phases. (D) Representative smFISH image used to determine cell area, with cell boundaries outlined by yellow dashed lines. (E) Probabilistic modeling on cell cycle phase transitions.

During different phases of cell growth, cells exhibit distinct metabolic behaviors [47, 48], which in turn influence the dynamics of mRNA-LNP delivery and expression. During the lag phase, cells acclimate to the new environment and do not immediately initiate division. Following adaptation, cells enter the early exponential phase, characterized by rapid proliferation and abundant nutrients [46], during which both mRNA-LNP uptake and protein synthesis occur at elevated rates. As nutrients become depleted, cells transition into the late exponential phase, where the LNP uptake rate begins to decline. This resource limitation is captured by a time-dependent growth rate modeled using Monod kinetics, as shown in Eq. (3). The substrate concentration *S*(*t*) is approximated by:

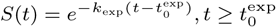

where *k*_exp_ is a time-scaling coefficient and 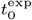 marks the onset of the exponential phase. The negative exponential form of *S*(*t*) reflects nutrient consumption over time. As cell growth ceases and viable cell density begins to decline, the mRNA-LNP delivery process correspondingly slows down.

### Cell Area-Dependent Division Model

While the macroscopic cell growth model in Eq. (3) captures the influence of culture conditions on population-level growth dynamics, accurately modeling the dilution of intracellular molecules during cell division requires a more detailed understanding of proliferation at the single-cell level. The cell cycle is typically divided into two main phases: G1-S and G2-M (see Figure 2A). For simplicity, the model excludes the possibility of cells entering the quiescent G0 phase. During the G1-S phase, cells grow by replicating DNA and synthesizing essential components such as proteins and ribosomes, leading to an increase in cell area. Once a critical size threshold is reached, cells transition into the G2-M phase [49]. During this phase, the rate of cell area expansion decelerates [43, 44, 45]. Upon completion of G2–M, the parent cell undergoes division, partitioning its genetic material and intracellular contents—including mRNAs and proteins—between two daughter cells.

The proposed cell area-dependent division model integrates three key data modules: (1) cell population growth curves derived from viability assays, (2) cell size distributions obtained via smFISH, and (3) the proportion of live cells in the G2-M phase, denoted by *γ*_G2-M_, measured through flow cytometry. The interdependence of these components is illustrated in Figure 2. To model cell cycle dynamics and transitions between the G1-S and G2-M phases, a Markov chain framework is employed, with the transition diagram shown in Figure 2A and the associated model components detailed in Figure 2E. At any time *t*_*h*_, let *z* (*t*_*h*_) denote the cell cycle stage of an individual cell, where *z* (*t*_*h*_) *∈* G1-S if the cell is in the G1-S phase, and *z* (*t*_*h*_) *∈* G2-M otherwise. The cell density growth rate, *µ*(*t*_*h*_), reflects the frequency of cell division and governs the transition rate from G2-M to G1-S, denoted by *r*_21_(*t*_*h*_), which determines the corresponding transition probability *P*_21_. Cell area growth primarily occurs during the G1-S phase. The transition probability from G1-S to G2-M, denoted by *P*_12_(*A*(*t*_*h*_) = *a*), is a function of the cell area *A*(*t*_*h*_). For a population of cells with area distribution density *q*(*A*(*t*_*h*_) = *a*), the expected transition probability is given by: *P*_12_ = ∫ *P*_12_(*a*)*q*(*a*)*da*. This formulation allows the model to account for heterogeneity in cell size across the population, linking individual cell growth dynamics to phase transition behavior in the cell cycle.

We describe the dynamics of cell area change during G1-S phase using an exponential growth model, which is commonly applied to both mammalian and plant cells [50, 51, 52]. Specifically, at any time *t*_*h*_, if *z*(*t*_*h*_) *∈* G1-S, the change in cell area *A*(*t*_*h*_) over the interval with length Δ*t* is given by,

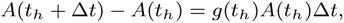

where *g*(*t*_*h*_) is the phase-dependent cell area growth rate. Notably, the growth rate during the G2-M phase is negligible compared to that observed in the G1-S phase.

The duration of the G2–M phase is modeled as an exponentially distributed random variable with rate parameter *r*_21_ (*t*_*h*_). Before re-entering the G1–S phase, the cell undergoes division, resulting in the partitioning of cell area and intracellular mRNA molecules between two daughter cells. As illustrated in Figure 1B, the area of one daughter cell is sampled from a binomial distribution:

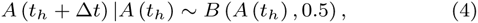

where *A*(*t*_*h*_) is the parent cell area at time *t*_*h*_ and *A*(*t*_*h*_ + Δ*t*) is the area of one daughter cell. According to mass balance, the area of the other daughter cell is *A*(*t*_*h*_) *− A*(*t*_*h*_ + Δ*t*). Molecules, including mRNA and proteins, are partitioned into daughter cells in proportion to their areas [28].

Given the macroscopic cell growth rate model as shown in Eq. (3), the transition rate *r*_21_ (*t*_*h*_) from G1-S to G2-M phase can be calculated as

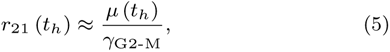

where *γ*_G2-M_ is the proportion of living cells in G2-M phase. The detailed derivation of Eq. (5) is presented in Supplementary Data Section 1.1. Using flow cytometry observations illustrated in Figure 2B, we can estimate the proportion of cells in G2-M phase, denoted by *γ*_G2-M_, by sorting cells in G1-S phase and G2-M phase. The stable and coordinated relationship between cell division and cell area growth is crucial for maintaining cell area homeostasis, ensuring that a cell population maintains a stable distribution of cell sizes (or areas) over time [51, 53]. This assumption is further supported by the results of the hypothesis test, as detailed in Supplementary Data Section 3.

Under this assumption, the transition probability from G1-S to G2-M phase for cell population can be derived as follows,

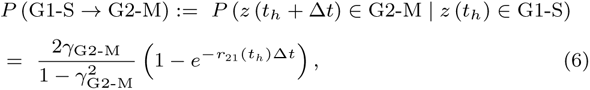

and *P* (G1-S *→* G2-M) is denoted as *P*_12_ for notational convenience. The derivation of Eq. (6) is detailed in Supplementary Data Section 1.2.

To investigate cell-to-cell variability and single-cell dynamics, we consider the conditional transition probability for a cell with area *a* to progress from the G1-S phase to the G2-M phase:

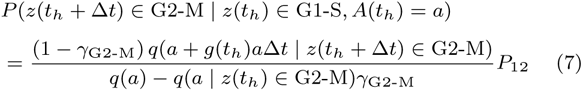

where *q*(*a*) represents the population distribution of cell area. The derivation of Eq. (7) is detailed in Supplementary Data Section 1.3. For any cell in the G2-M phase, the conditional distribution of cell size can be derived using Bayes’ rule, i.e.,

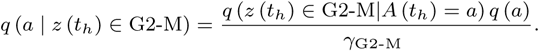

Following the study [54], the conditional probability that a cell is in the G2-M phase, given that its area is *a*, is modeled using a sigmoid function,

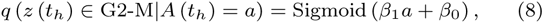

with *β*_1_ *>* 0, where *β*_1_ and *β*_0_ are the linear transformation coefficients of cell area. Then, by substituting Eq. (8) into Eq. (7), we can obtain the conditional transition probability from G1-S to G2-M phase as,

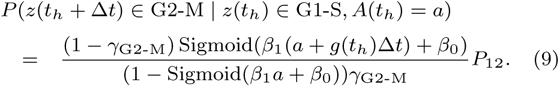

The detailed derivation is provided in Supplementary Data Section 1.3.

### Stochastic mRNA-LNP Uptake

The mRNA–LNP uptake process involves two key steps: binding and endocytosis. The uptake rate of the mRNA-LNP, denoted by *v*_up_(*t*_*h*_), is governed by the size of the nanoparticle. Assuming a spherical nanoparticle and a relatively flat cell membrane, the mRNA–LNP complex is enveloped by the membrane through receptor–ligand interactions. During endocytosis, free receptors on the membrane diffuse toward the LNP and bind to ligands on its surface. Once all ligands are engaged, the complex is fully wrapped and internalized by the cell (see Figure 1). The size of the LNP and its ligand density determine the number of available binding sites, which in turn dictates the number of binding events required for successful uptake.

Given the nanoparticle size *s*_LNP.size_(*t*_*h*_), the membrane wrapping time is calculated according to the model proposed by Gao et al. [5] as follows,

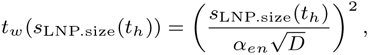

where *D* is diffusivity of membrane receptors, and *α*_*en*_ is a constant referred to as the ‘speed factor’, derived from the energy balance equation:

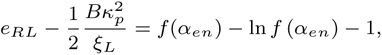

with 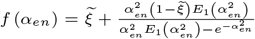. Here *e*_*RL*_ is the receptor-ligand adhesion energy, estimated to be approximately 15*k*_*B*_*T* at 300 K [55], a representative temperature for nanoparticle delivery studies [5]. The bending modulus *B* is typically taken as 20 *k*_*B*_*T* [55, 56], and the nanoparticle curvature is given by 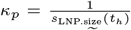. Ligand density on the LNP surface is denoted by *ξ*_*L*_, and *ξ* represents the ratio of receptor to ligand density; both are model-fitted parameters. For computational simplicity, *α*_*en*_ is treated as a constant, determined using the average LNP size. Since mRNA loading is proportional to the LNP volume [18, 19], the internalization rate of mRNA molecules is given by:

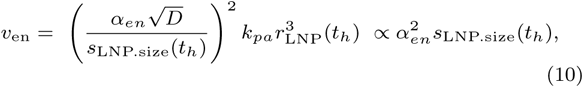

where *k*_*pa*_ is the proportionality constant relating mRNA loading to LNP volume. Thus, the rate of mRNA endocytosis is governed by both energy dynamics (*α*_*en*_) and volume-based LNP loading.

Since mRNA–LNP uptake occurs via ligand–receptor binding, the extracellular mRNA concentration directly influences intracellular mRNA levels [5, 24, 57]. To account for cell–LNP interactions, we define a normalized extracellular LNP concentration as:

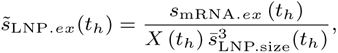

where 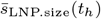 is the average LNP radius and *X*(*t*_*h*_) is the total cell count at time *t*. The term 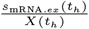 represents the mRNA concentration per cell, which is further scaled by the LNP volume 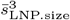 (*t*_*h*_) to reflect the effective concentration of LNPs available for uptake.

The mRNA uptake rate *v*_up_(*t*_*h*_) is modeled using an empirical Monod-type formulation that combines LNP binding and internalization dynamics:

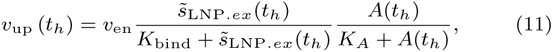

where *v*_en_ is the LNP size-dependent internalization factor, as defined in Eq. (10). The first saturation term in Eq. (11) captures the dependence of uptake rate on the normalized extracellular mRNA–LNP concentration 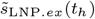, with *K*_bind_ representing the half-saturation constant for ligand–receptor binding. This constant may be influenced by factors such as LNP surface charge and cell membrane properties [6]. The second saturation term accounts for the effect of cell surface area *A*(*t*_*h*_) on uptake rate. Experimental studies [7, 58] and observations (see Figure 3D) indicate that larger cell size enhances uptake. The parameter *K*_*A*_ denotes the half-saturation constant for area-dependent effects, with saturation reflecting biological constraints such as limited energetic capacity for endocytosis or receptor-mediated regulation [59, 60].

**Fig. 3:**
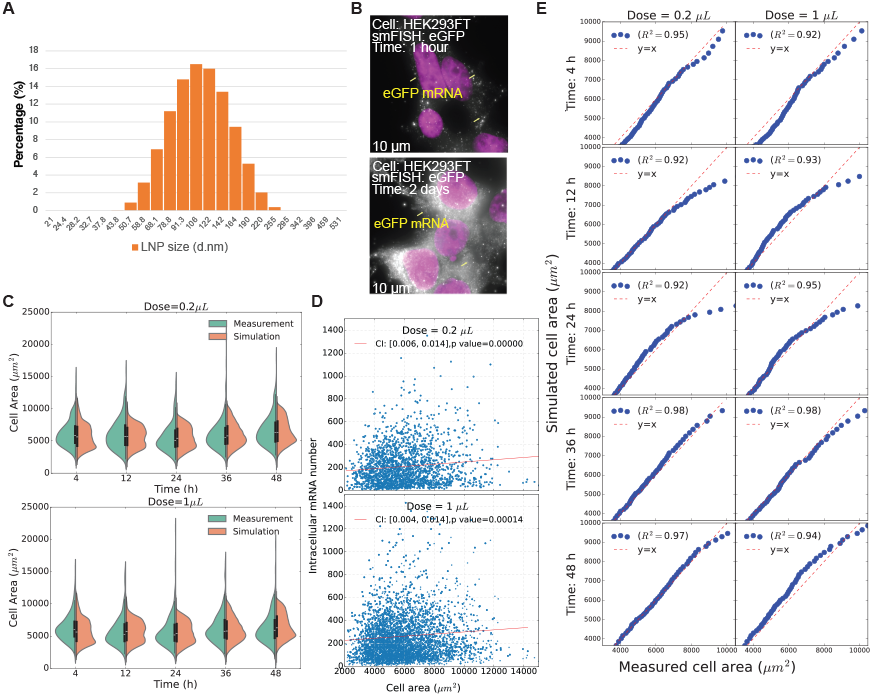
Measurements and predictions on cell area dynamics and LNP size distribution. (A) Histogram of experimentally measured LNP size distribution. (B) Representative smFISH imagings showing intracellular distribution of eGFP mRNA at 1 hour and 2 days post-transfection in HEK293FT cells. (C) Violin plots comparing observed and simulated cell area distributions over time for two administered doses: 0.2 *µL* and 1*µL*. (D) Scatter plot analysis reveals a weak positive correlation between cell area and intracellular mRNA copy number, supported by statistically significant two-sided p-values. (E) Q-Q plots comparing simulated and observed cell area distributions over time.

### mRNA Degradation and Exocytosis

To account for mRNA degradation and exocytosis, the intracellular mRNA concentration dynamics are modeled as follows:

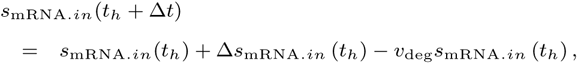

where *v*_deg_ denotes the intracellular mRNA degradation rate. The change in intracellular mRNA concentration due to uptake and exocytosis is given by:

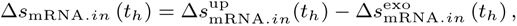

where the change due to mRNA uptake is modeled as a normally distributed random variable: 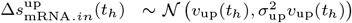 with *v*_up_(*t*_*h*_) obtained from Eq. (11) and 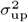 representing the uptake variability factor.

Additionally, mRNA exocytosis is modeled using empirical Monod kinetics, incorporating stochastic variability through a normally distributed random process:

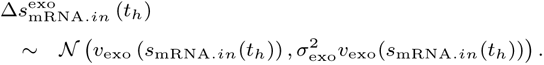

Here, the mean exocytosis rate is defined by a Monod-type saturation function:

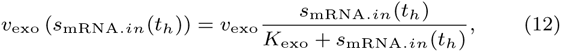

where *v*_exo_ is the maximum mRNA efflux rate, *K*_exo_ is the empirical half-saturation constant characterizing mRNA efflux, and 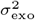 represents the coefficient of variation, capturing stochastic fluctuations in the exocytosis process. This formulation captures both the nonlinear saturation behavior of mRNA efflux and its inherent variability, enabling more realistic simulation of intracellular transport dynamics.

Accordingly, since naked RNAs released via exocytosis cannot re-enter cells due to electrostatic repulsion, the extracellular mRNA concentration—encapsulated within LNPs—is updated by accounting for uptake dynamics across all viable cells with density *X*(*t*_*h*_), as follows:

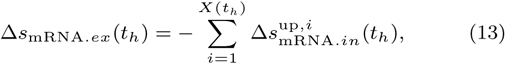

where 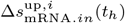 denotes the mRNA uptake rate by the *i*-th cell at time *t*_*h*_. This formulation captures the net depletion of extracellular mRNA due to cellular uptake, excluding contributions from exocytosed naked mRNA.

### mRNA Translation

The above model describes the process from mRNA–LNP uptake to intracellular mRNA release. Once cells adapt to their environment, they exhibit distinct translation rates across different metabolic phases. The protein accumulation in a single cell, denoted by Δ*s*_Protein_ (*t*_*h*_), is modeled as a normally distributed random variable:

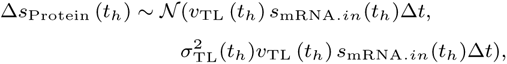

where *v*_*T L*_ (*t*_*h*_) is the phase-specific translation rate, and 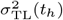 is the translation variability coefficient. This variability is influenced by stochastic factors such as mRNA stability and innate immune activation [9].

#### Model Inference Method

The proposed MIMS model, built upon a single-cell kinetic framework, enables simulation of gene and protein expression trajectories at the single-cell level. The metabolic phase-specific cell growth model is initially calibrated using macroscopic measurements of viable cell density. In parallel, the cell division model is calibrated based on the estimated growth rate and the observed distribution of cell areas. Given the inputs—including the initial extracellular mRNA concentration *s*_mRNA.*ex*_(0), the calibrated single-cell division model, and the initial LNP size distribution *s*_LNP.size_(0)—MIMS simulates the distribution of multivariate state trajectories ***τ*** = {***u***(*t*_0_), ***u***(*t*_1_), …, ***u***(*t*_*H*_)} according to the following probabilistic formulation:

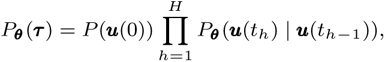

where the state transition model *P*_***θ***_(***u***(*t*_*h*_)|***u***(*t*_*h−*1_)) is governed by mechanistic parameters ***θ***. These parameters define modular processes including cell growth, area-dependent division, mRNA–LNP uptake, mRNA degradation, exocytosis, and translation.

The multi-scale measurements used in this study provide heterogeneous and partial observations of the underlying state variables. These measurements include: (1) initial measurements of extracellular mRNA concentration and LNP size distribution, obtained using qRT-PCR and DLS, respectively; (2) time-resolved single-cell joint distributions of intracellular mRNA counts and cell sizes obtained via smFISH; and (3) time-resolved single-cell distributions of protein expression levels obtained via flow cytometry. For the *k*-th potency measurement, denoted by *s*_*k*_ (i.e., intracellular mRNA count or protein expression), collected at time *t*_*k,i*_ with time index *i* = 1, 2, …, *H*_*k*_, the empirical distribution of single-cell expression is represented by *P* (*s*_*k*_(*t*_*k,i*_)). The model-predicted distribution is denoted by 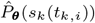. To optimize ***θ***, we minimize the Wasserstein loss [61], which quantifies the distance between the empirical and predicted distributions:

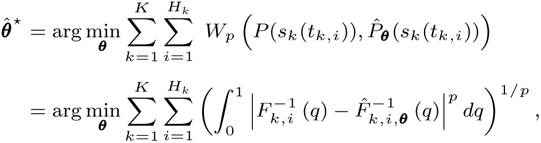

where *K* denotes the number of potency measurements, 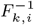 and 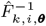 are the quantile functions of the empirical and predicted distributions, respectively. In our analysis, we let *p* = 1, simplifying the Wasserstein distance to the average absolute difference between empirical quantiles: 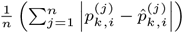, where 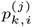 and 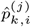 are the *j*-th empirical quantile of *P* (*s*_*k*_(*t*_*k,i*_)) and 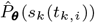, respectively, and *n* is the number of quantiles used.

To ensure the Wasserstein distance is comparable across datasets with varying scales, we define the loss function using a scaled version of the Wasserstein distance:

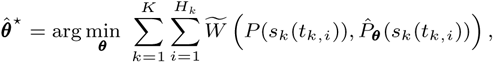

where the scaled Wasserstein distance is given by:

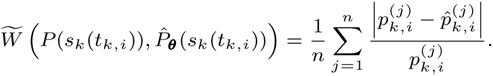

The scaling ensures that the loss remains consistent regardless of the magnitude of the underlying data distributions. To optimize the model parameters, we employ the Adam optimizer [62].

Therefore, the developed cell-based potency assessment framework integrates mechanistic insights with heterogeneous experimental datasets collected across multiple scales, using model parameter optimization. By incorporating iterative updates informed by advanced and cost-effective assays, this approach enhances our understanding of the dynamics and variability of mRNA–LNP delivery and expression at the cellular level.

## Results

### In-Vitro Assays and Data Analytics

This section presents the experimental observations from the *in vitro* studies on eGFP mRNA-LNP delivery and expression, conducted using HEK293FT cells over a 7-day period. At the macroscopic scale, qRT-PCR was employed to measure the initial extracellular mRNA concentration. Concurrently, cell viability data were collected to characterize the metabolic phases of cell growth. At the single-cell scale, flow cytometry was used to quantify the proportion of viable cells across different stages of the cell cycle. smFISH assays provided time-course measurements of intracellular eGFP mRNA copy numbers, while brightfield imaging of fixed cells was used to determine cell area. Additionally, eGFP protein expression levels were assessed via flow cytometry. To investigate the impact of nanoparticle size on cellular uptake and potency variability, the size distribution of mRNA-LNPs was measured using DLS [20].

### Cell Culture

HEK293FT cells (Thermo Fisher cat# R700-07) were cultured following the manufacturer’s protocol and maintained in high glucose Dulbecco’s modified Eagle’s medium (DMEM) with GlutaMax (Fisher Scientific cat# 10566016), 10% fetal bovine serum (FBS), 1% MEM non-essential amino acids (Thermo Fisher cat# R70007), 1% supplementary GlutaMax (Fisher Scientific cat# 35-050-061), 1% MEM sodium pyruvate (Thermo Fisher cat# 11360070), 1% Pen-Strep (Fisher Scientific cat# BW17-602E), and 500 *µg*/mL geneticin (Thermo Fisher cat# 10131035). Cell lines were tested PCR-negative for mycoplasma contamination (ATCC cat# 30 1012K).

#### Viable Cell Number

To test the viable cell number, a total of 12,500 HEK293FT cells were seeded into each well of a 96-well ELISA assay microplate (Fisher Scientific cat# 07-000-627) in 150 *µL* of growth medium and incubated for 24 hours to allow cell attachment. Cell number and viability were assessed at 1, 4, 12, 24, 48, 72, 96, 120, 144, and 168 hour (h) using a TC20™ Automated Cell Counter (Bio-Rad), following the manufacturer’s protocol. Trypan blue exclusion (Bio-Rad, #145-0022) was used to distinguish viable cells from non-viable ones.

The cell viability measurements from three biological replicates are presented in Figure 2C. During the initial 24 hours, cell viability increases gradually, reflecting a slow proliferation rate. After this period, a sharp rise in both the expected cell count and inter-replicate variability is observed. Between 120 and 144 hours, the cell count begins to decline, indicating a reduction in viable cells following the peak growth phase. Based on the observed growth kinetics, three distinct metabolic phases of cell growth are identified: ➀ Lag phase (0–3 hours): Characterized by minimal cell division as cells adapt to the culture conditions. ➁ Exponential phase (4–95 hours): Marked by rapid cell proliferation. This phase can be further subdivided into early exponential phase (4–23 hours) and late exponential phase (24–95 hours). ➂ Decline phase (96–168 hours): Cell proliferation decelerates and subsequently diminishes, likely due to nutrient exhaustion, accumulation of cytotoxic metabolites, or other stress-induced inhibitory factors within the culture environment.

#### Cell Cycle

Cells in the G1-S and G2-M phases were sorted as shown in Figure 2B. A 1 mL suspension of HEK293FT cells was prepared at a concentration of 1 *×* 10^6^ cells/mL in growth medium. Vybrant™ DyeCycle™ Violet (Thermo Fisher Scientific, Cat# V35003) was added to a final concentration of 5 *µM* (1 *µL* per 1 mL of cell suspension), mixed gently, and incubated at 37^*°*^C for 30 min, protected from light. Cells were maintained at 37^*°*^C until flow cytometry acquisition and analyzed live—without washing or fixation—using 405 nm excitation laser and emission detection centered at ~450/50 nm.

Among the 31, 407 cells analyzed, 21, 932 cells (69.8%) were classified as being in the G1-S phase, while 9, 475 cells (30.2%) were in the G2-M phase. Accordingly, the proportion of viable cells in the G2-M phase is assumed to be constant and set as *γ*_G2-M_ = 30.2% in Eq. (5).

#### Initial Extracellular mRNA Copy Number

qRT-PCR was employed to quantify the initial mRNA copy numbers of mRNA-LNPs in the extracellular medium. Measurements from three biological replicates are summarized in Table 2. LNP-encapsulated RNA was extracted using Trizol reagent (Thermo Scientific cat# 15-596-026) and resuspended in nuclease-free water. Residual genomic DNA was removed by treatment with DNase I (Thermo Scientific cat# PI89836). Reverse transcription of RNA was performed using SuperScript III Reverse Transcriptase (Thermo Scientific cat# 18-080-093). Quantification of the resulting cDNA was carried out using Luna Universal qPCR Master Mix (New England Biolabs cat# M3004S). The sequences are provided in Supplementary Data Section 2.

**Table 2.** Initial extracellular eGFP mRNA molecule copy numbers measured by qRT-PCR.

| | Dose = 0.2 μL | Dose = 1 μL |
| --- | --- | --- |
| **Replicate 1** | 109, 500, 488 | 598, 183, 138 |
| **Replicate 2** | 41, 794, 286 | 445, 503, 342 |
| **Replicate 3** | 48, 603, 399 | 201, 679, 148 |
| **Average** | 66, 632, 724 | 415, 121, 876 |
| **Standard deviation** | 37,280,355 | 199, 990, 321 |

#### Intracellular mRNA Copy Number and Cell Area

The smFISH assay was employed to obtain time-course measurements of single-cell mRNA copy number and corresponding cell area. smFISH enables visualization of individual RNA molecules as discrete fluorescent spots within cells. By acquiring image stacks, the spatial distribution of each RNA species can be accurately reconstructed at the single-cell level. A total of 12,500 HEK293FT cells were seeded into each well of an 8-well chambered coverglass (Thermo Scientific cat# 12-565-470)) in 300 *µL* of growth medium and incubated for 24 hours to allow cell attachment. Cells were treated with two volumes (0.2 *µL* and 1 *µL*) of eGFP mRNA-loaded lipid nanoparticles (LNPs) at various time points: 48 h, 36 h, 24 h, 12 h, 4 h, and 1 h prior to fixation. Cells were fixed with 4% formaldehyde solution and permeabilized overnight in 70% ethanol. For smFISH, 1 *µL* of eGFP-specific probe mixture was added to 50 *µL* hybridization buffer (containing 10% dextran sulfate, 10% formamide, and 2x SSC) and the samples were incubated overnight at 37°C. Following hybridization, samples were washed twice with wash buffer (10% formamide and 2x SSC), and cell nuclei were stained with DAPI (Thermo Scientific cat# EN62248). Imaging was performed using a Nikon inverted research microscope equipped with a Plan Apo *λ* 100×/1.45 oil immersion objective. Four wavelengths were recorded: Alexa 488, Alexa 594, DAPI, and Brightfield. The smFISH images were processed using a custom analysis pipeline to segment individual cells, apply spot detection thresholds, and count RNA spots. Single-cell segmentation was conducted using the MATLAB-based RajLab Image Tools suite, available at https://bitbucket.org/arjunrajlaboratory/rajlabimagetools/wiki/Home. The probe sequences are provided in Supplementary Data Section 2.

Experiments were conducted in three biological replicates using 0.2 *µL* and 1 *µL* doses, with 100 cells sampled at each time point for each dose and replicate. Figure 3D presents a scatter plot of cell area versus intracellular mRNA count, revealing a weak positive correlation under both dosing conditions with significant two-sided p-value. This relationship is substantially weaker than the linear proportionality reported by Rees et al. [7], suggesting that mRNA-LNP binding and uptake could be modulated by saturation and inhibition mechanisms intrinsic to living cells—such as energy availability and receptor expression.

To characterize the single-cell distribution of intracellular mRNA levels, smFISH data from all biological replicates were pooled and visualized as green violin plots in Figure 5A. Summary statistics corresponding to these distributions are provided in Table 3. Outliers were removed using the filtration procedure described by Tukey [63] to enhance model robustness. The experimental results align with previous studies [12, 57], showing that intracellular mRNA levels rise sharply within the first 24 hours and subsequently plateau, maintaining a relatively stable concentration over the remainder of the observation period.

**Table 3.** Statistics of intracellular mRNA number measured by smFISH,.

| Time (h) | Dose = 0.2 μL | | Dose = 1 μL | |
| --- | --- | --- | --- | --- |
|  | Mean | Std | Mean | Std |
| **1** | 49.13 | 25.85 | 93.69 | 36.14 |
| **4** | 107.04 | 50.94 | 133.71 | 57.37 |
| **12** | 174.76 | 102.53 | 206.28 | 118.35 |
| **24** | 223.66 | 133.26 | 301.39 | 158.67 |
| **36** | 281.41 | 154.46 | 302.76 | 136.85 |
| **48** | 259.11 | 152.99 | 319.18 | 158.58 |

To evaluate the distribution of cell area over time, we constructed Q–Q plots to assess the temporal consistency of cell area distributions, as described in Supplementary Data Section 3. The results indicate that the distribution stabilizes after 4 hours, as illustrated in Figure 3C.

#### eGFP Intensity

Flow cytometry was used to quantify eGFP protein expression intensity across three biological replicates, using 0.2 *µL* and 1 *µL* doses (see Figure 4). A total of 12,500 HEK293FT cells were seeded in each 96-well ELISA Assay Microplates (Fisher Scientific cat# 07-000-627) with 150 *µL* of growth medium and incubated for 24 hours to allow cell attachment. Cells were treated with two volumes (0.2 *µL* and 1 *µL*) of eGFP mRNA-loaded lipid nanoparticles (ProMab cat# PM-LNP-0021) across three rows of wells at the following time points: 168 h, 144 h, 120 h, 96 h, 72 h, 48 h, 24 h, 12 h, 4 h, and 1 h prior to analysis. Following treatment, the medium was aspirated, and cells were washed once with phosphate-buffered saline (PBS) containing 5% bovine serum albumin (BSA). Cells were then resuspended in 200 *µL* PBS with 5% BSA for flow cytometry analysis. Flow cytometry was performed using a Beckman Coulter Cytoflex Cell Analyzer, with fluorescence detection in the KO525 channel. Data were preprocessed and analyzed for event counts using FlowJo software. When comparing eGFP-expressing cells to the negative control, we found that cellular autofluorescence significantly interferes with the detection of fluorescence signals in cells exhibiting low eGFP expression [64]. This interference introduces considerable uncertainty in the quantification of low-intensity eGFP signals. To ensure both interpretability and robustness, our analysis is restricted to high-expression cells, defined as those exhibiting eGFP intensity values *≥* 360 RFU. This threshold corresponds to the initial protein intensity cutoff, as indicated by the vertical line in Figure 4.

**Fig. 4:**
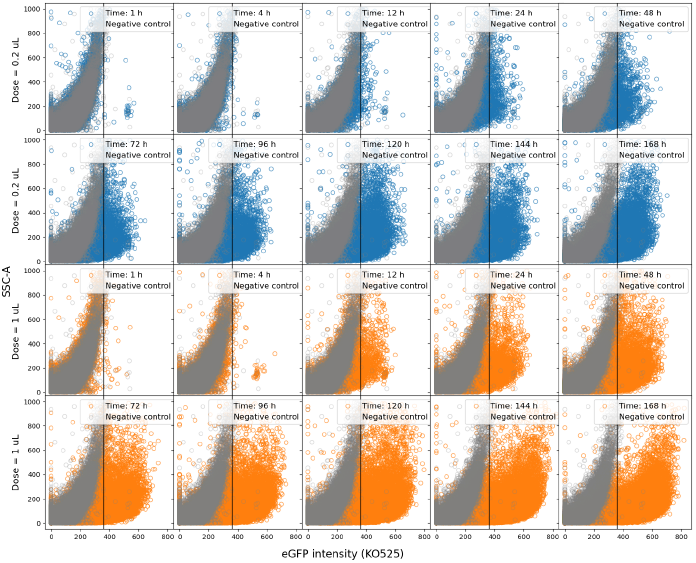
Scatter plots illustrate the time-course measurements of eGFP expression. Each row corresponds to a different dose (0.2*µL* and 1*µL*), while columns correspond to time points post-treatment. The horizontal axis denotes the fluorescence intensity measured in the KO525 channel. The vertical axis indicates side-scattered light area (SSC-A). Blue and orange points denote experimental samples treated with 0.2*µL* and 1*µL* doses, respectively. Gray points represent negative controls, which is measured at 0 hours post-treatment.

**Fig. 5:**
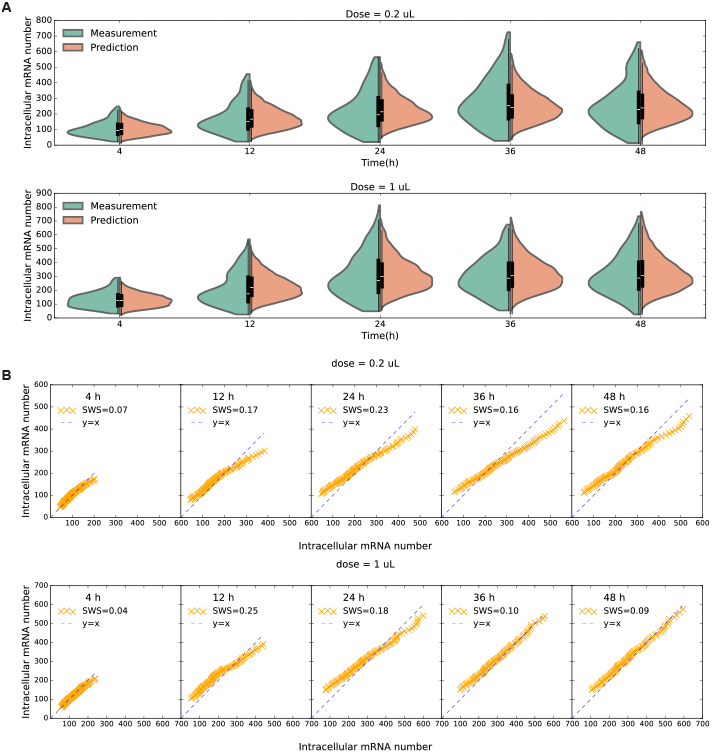
Predictive results of intracellular mRNA copy number per cell. The predictive results demonstrate the model’s performance when trained on data from one dose condition and validated on another. (A) Violin plots illustrate both measured and predicted intracellular mRNA copy numbers over time for two dose conditions: 0.2*µL* (top panel) and 1*µL* (bottom panel). Experimental measurements are shown in green, while model predictions are shown in orange. (B) Quantile–Quantile (Q–Q) plots compare predicted and measured intracellular mRNA copy numbers at selected time points (4–48 hours) for the same two dose conditions: 0.2*µL* (top panel) and 1*µL* (bottom panel).

Then, high-intensity data from all three replicates were pooled and visualized as green violin plots in Figure 6A, whose statistics are presented in Table 4. Outliers were removed using Tukey’s filtration method [63]. For both 0.2*µL* and 1*µL* doses of eGFP mRNA-LNPs, the mean and variance of eGFP expression increased gradually over time, reaching a plateau at approximately 120 hour. Beyond this point, the protein expression level remains stable, suggesting convergence to a translation saturation threshold.

**Table 4.**
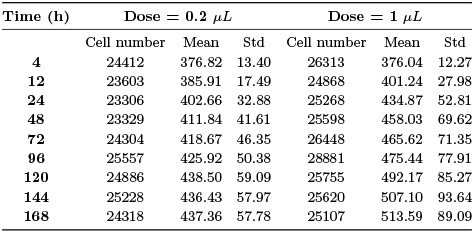
Mean and standard deviation (Std) of eGFP expression intensity (RFU) for protein expression *≥* 360 RFU.

**Fig. 6:**
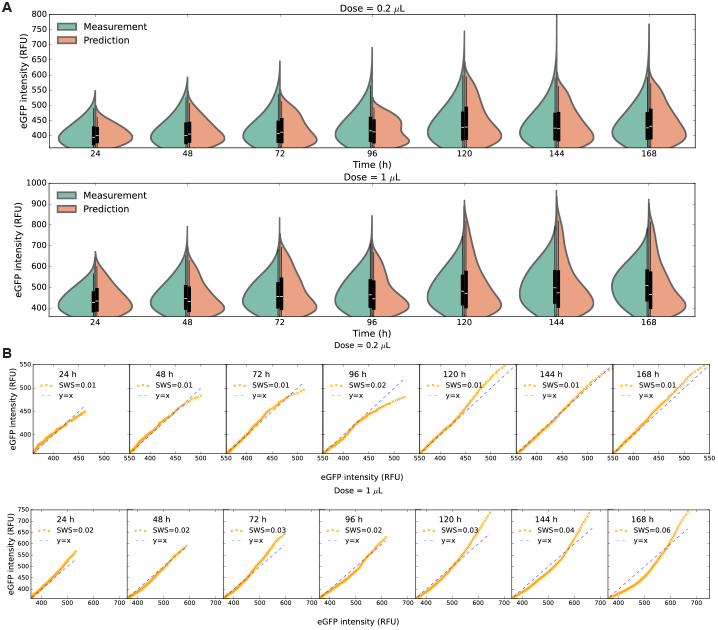
Predictive results of protein expression level concentration. (A) Violin plots illustrate the distribution of both measured and predicted positive eGFP intensity (RFU *≥* 360) over time under two dosing conditions: 0.2*µL* (top panel) and 1*µL* (bottom panel). Green curves represent experimental measurements, while orange curves indicate model predictions. (B) Quantile–Quantile (Q–Q) plots compare predicted versus observed eGFP expression levels at selected time points (24–168 hours). Data points include both fitting results (yellow crosses) and validation results (purple circles) for the two dosing conditions: 0.2*µL* (top panel) and 1*µL* (bottom panel).

#### LNP Size

Dynamic Light Scattering (DLS) was used to characterize the particle size distribution of mRNA-loaded lipid nanoparticles (LNPs) in phosphate-buffered saline (PBS). For measurements in PBS, LNPs were diluted at a ratio of 1:500. DLS detects fluctuations in scattered light intensity caused by Brownian motion of particles, allowing for the determination of their size distribution. This technique provides insight into the physical properties of the nanoparticles under different environmental conditions. For further details, refer to [20].

The size distribution of lipid nanoparticles (LNPs) is presented in Figure 3A. The overall particle diameters distribution closely approximates a normal distribution, with mean value of 116.42 nm and standard deviation of 38.12 nm.

### MIMS Model Implementation and Validation

To characterize cellular mechanisms, we first calibrated the cell density model described in Eq. (3) using cell viability measurements. This was followed by fitting the cell division model to time-course data of cell area. We then assessed the predictive accuracy of the multi-scale kinetic MIMS model by comparing its outputs against intracellular mRNA copy numbers and eGFP protein expression levels. Upon successful validation, we performed *in silico* experiments to explore how dosage, LNP size, and cell area influence the dynamics and variability of gene and protein expression. The simulation results show strong agreement with previously reported findings, further demonstrating the reliability and robustness of the proposed MIMS framework.

### Simulation of Cell Growth and Cell Division

HEK293FT cells are widely employed in mRNA-LNP delivery studies due to their high transfection efficiency [65, 66], robust protein expression capabilities [67, 68], and rapid proliferation rate [26]. Our fitted cell growth model accurately reproduces the experimentally measured HEK293FT cell density over time, as shown in Figure 2C. Importantly, the model captures all three phases of cell growth, aligning well with observations from batch-cultured HEK293FT systems [26, 69, 70].

Building on this validated macroscopic model of cell population dynamics, we developed a complementary cell division model that accurately predicts the distribution of cell areas, as shown in Figure 3C. Together, these models offer distinct yet synergistic perspectives on cell proliferation. The cell growth model describes population-level dynamics driven by external resource availability, while the cell area-dependent division model provides a finer-grained view at the single-cell level, incorporating internal regulatory factors such as cell size and cell cycle progression. During proliferation, cells tend to maintain a stable distribution of cell areas, indicating that cell size plays a regulatory role in division [51, 71, 53, 72]. Assuming a constant proportion of cells across different cell cycle phases, both the division rate and the transition rate from the G1-S phase to the G2-M phase are modulated by the resource-dependent cell growth rate, as described in Eq. (5) and Eq. (6). Notably, larger cells in the G1-S phase exhibit a higher probability of progressing to the G2-M phase, as modeled by Eq. (7).

### Model Prediction of mRNA-LNP Potency

To evaluate the predictive performance of the MIMS model in capturing single-cell gene and protein expression dynamics and variability, we employed cross-validation. Specifically, the model was trained on one dose dataset and independently validated against a separate dose dataset to ensure robustness. For protein expression intensity prediction, we focused on values exceeding the negative control threshold (360 RFU), as illustrated in Figure 4. This threshold was established based on observations from the negative control to mitigate the confounding effects of cell autofluorescence [13, 8]. Since too few cells positively express eGFP before 24 hours when dose is 0.2*µL*, we only predict the protein expression after 24 hours.

Figure 5 visualizes the predicted distribution of intracellular mRNA copy numbers at the single-cell level. Quantitative evaluation of the model’s performance—including relative errors in the mean (*µ*), standard deviation (*Σ*), and the overall distributions of gene and protein expression—is summarized in Table 5. Violin plots illustrate the empirical distributions from both model predictions and experimental assay measurements, while Quantile-Quantile (QQ) plots reveal the linear correspondence between predicted and observed distributions. The proposed multi-scale kinetic model demonstrates strong and consistent predictive performance in capturing gene and protein expression dynamics, as well as cell-to-cell variability. Furthermore, the model generalizes well to dose-response conditions not included in the training data, providing accurate and reliable predictions beyond the scope of the original experiments.

**Table 5.** Model prediction relative errors of the mean *µ*, standard deviation *Σ*, and overall distribution of cell-based mRNA-LNP potency.

| Dose ($\mu\text{L}$) | Intracellular mRNA concentration | | | Protein expression level | | |
| --- | --- | --- | --- | --- | --- | --- |
| | $\frac{\vert \hat{\mu}-\mu\vert }{\mu}$ | $\frac{\vert \hat{\sigma}-\sigma\vert }{\sigma}$ | Scaled Wasserstein distance | $\frac{\vert \hat{\mu}-\mu\vert }{\mu}$ | $\frac{\vert \hat{\sigma}-\sigma\vert }{\sigma}$ | Scaled Wasserstein distance |
| 0.2 | 0.03 | 0.29 | 0.16 | 0.01 | 0.10 | 0.01 |
| 1 | 0.06 | 0.19 | 0.13 | 0.02 | 0.21 | 0.03 |

As shown in Figure 5A, smFISH assay results reveal that intracellular mRNA concentrations increase rapidly within the first 24 hours post-treatment and subsequently stabilize at a steady-state level. While Ashraf *et al*. [57] proposed a fixed saturation threshold for single-cell mRNA expression, our experimental data suggest that the saturation level is dose-dependent. Specifically, at a dose of 0.2 *µL*, the mRNA copy number plateaus around 260 spots per cell, whereas at a dose of 1 *µL*, the saturation level increases to approximately 300 spots per cell.

The developed multi-scale kinetic model accounts for the observed differences in saturation levels and links them to the dose-dependent uptake rate of mRNA-LNPs. Specifically, the LNP uptake rate increases with dose, as described by Eq. (11), until it reaches a saturation state. As exocytosis transports some mRNA back into the extracellular environment, the exocytosis rate is constrained by a maximum rate *v*_exo_, as defined in Eq. (12). At a fixed dose level, individual cells regulate intracellular mRNA concentration through a dynamic balance of three processes: influx of mRNA via LNP uptake, loss of mRNA through exocytosis (the cellular mechanism for releasing molecules), and dilution of mRNA during cell division. This regulatory mechanism enables cells to maintain mRNA homeostasis despite external perturbations and growth. Cell size further influences this balance. Larger cells tend to exhibit faster mRNA-LNP uptake and are more prone to division, while smaller cells show slower uptake and exocytosis rates and are less likely to divide. These size-dependent dynamics add another layer of complexity to intracellular mRNA regulation.

### Simulation-Based Prediction of Potency Across Varying Doses

The validated multi-scale kinetic model was used to perform *in silico* experiments to investigate the effect of dose on mRNA-LNP potency. To characterize the dose–response relationship, simulations were conducted at five dose levels: 0.1 *µL*, 0.2 *µL*, 0.5 *µL*, 1.0 *µL*, and 2.0 *µL*, with results shown in Figure 7. In each case, the initial mRNA concentration was scaled proportionally to the dose, assuming encapsulation within mRNA-LNPs of uniform size, as illustrated in Figure 3A. The resulting dose–response curve exhibits a sigmoid-like profile, consistent with observations by Yasar et al. [13]. As the dose increases, the variability in intracellular mRNA concentration also rises, aligning with findings from Rees et al. [7]. Notably, the MIMS model, unlike the live-cell imaging approach used by Leonhardt et al. [12], captures single-cell dynamics using cost-efficient fixed-cell imaging, enabling reliable dose–response predictions without the need for live-cell imaging techniques.

**Fig. 7:**
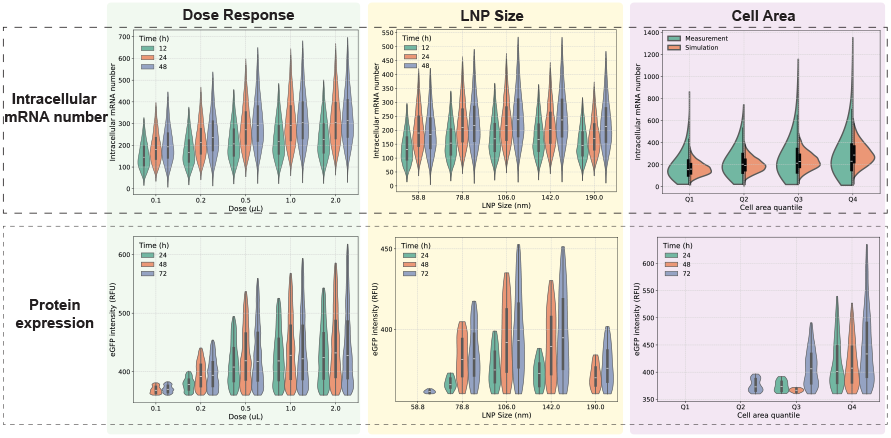
*In silico* experiment results characterizing the impact of dose, LNP size, and cell area on mRNA-LNP delivery outcomes. Violin plots of predicted intracellular mRNA copy number (top row) and eGFP protein expression (bottom row) across three sources of variability: dose (left), LNP size (middle), and cell area (right). For each condition, distributions are shown at selected time points (mRNA: 12 h, 24 h, 48 h; protein: 24 h, 48 h, 72 h). In the top row of cell area panel, green and yellow denote experimental observations and model simulations, respectively. For LNP size and cell area *in silico* experiments, the mRNA-LNP doses are 0.2 *µL*.

The simulation results (Figure 7) indicate a sharp increase in mRNA uptake efficiency with increasing dose, reaching a plateau at approximately 0.5 *µL*. This trend is consistent with experimental findings reported by Leonhardt et al. [12], Rees et al. [7], and Jiang et al. [73]. Given comparable eGFP nanoparticle sizes, the model-generated dose–response curve closely matches that of Leonhardt et al., despite relying solely on fixed-cell assay data. Additionally, the simulations reveal that cell-to-cell variability in intracellular mRNA levels increases with dose, further corroborating the findings of Rees et al.. The predicted protein expression response also aligns well with trends and variability reported in the literature, such as studies by Yasar et al. [13], Shen et al. [74], and Munson et al. [25].

Notably, compared to previous studies, the developed multi-scale kinetic model offers a more interpretable framework for predicting dose–response behavior. Since the uptake rate in Eq. (11) follows the Monod equation, the mRNA-LNP internalization rate is constrained by the parameter *v*_en_, which is influenced by factors such as LNP size, ligand density, and receptor density. When cell area and LNP size are held constant, increasing the dose enhances the ligand–receptor binding rate. This binding rate is modulated by the half-saturation constant *K*_bind_, which reflects underlying cellular regulatory mechanisms, including energy availability and cell culture conditions. Consequently, within the proposed framework, the selection of the mRNA-LNP uptake rate is inherently linked to the cell growth rate, providing mechanistic insight into dose-dependent variability.

### LNP Size-Dependent Potency in mRNA-LNP Delivery

The validated multi-scale kinetic model was employed to perform *in-sillico* experiments investigating the influence of LNP size on delivery potency. Specifically, we simulated the potency of mRNA-LNP formulations across a range of LNP sizes to assess their impact on the delivery system’s effectiveness. Five representative LNP sizes were selected based on quantiles of the experimental size distribution: 58.8 nm (4 quantile), 78.8 nm (22 quantile), 106 nm (53 quantile), 142 nm (83 quantile) and 190 nm (98 quantile). To simplify the analysis, size variation within each case was neglected. The initial dose for all simulations was set to 0.2 *µL*.

As illustrated in Figure 7, the simulated average potency reaches its peak when the LNP size is approximately 106-142 nm. Deviations from this optimal size—either smaller or larger—result in reduced potency. This finding is consistent with observations reported in the literature from both *in vitro* even *in vivo* studies [75]. Within the optimal size range, cell-to-cell variation in potency is maximized, suggesting a highly heterogeneous response across the cell population. In contrast, smaller LNPs exhibit limited mRNA encapsulation capacity.

Although they demonstrate higher binding rates at a fixed dose, their overall mRNA uptake is inefficient due to the reduced loading per particle. Additionally, the increased membrane curvature induced by smaller LNPs elevates the elastic energy cost during internalization [5]. Conversely, excessively large LNPs suffer from lower binding rates and prolonged internalization times. In both suboptimal size regimes, mRNA uptake becomes uniformly low across the cell population, leading to diminished variability in cellular response.

We conducted a systematic simulation study to construct a response surface that characterizes the influence of LNP size on the expected intracellular mRNA copy number. This surface serves as a guide for selecting the optimal LNP size to maximize mRNA delivery efficiency. The simulated response surface, which exhibits a quadratic-like trend (Figure 8), enables identification of an optimal size range based on average mRNA uptake. Using intracellular mRNA copy numbers measured at 24 and 48 hours as evaluation criteria, we performed an exhaustive search and identified an optimal LNP size range of 106–142 nm, with a peak around 110 nm. This finding is consistent with experimental observations reported in the literature. For instance, Walther et al. [22] reported an optimal LNP size of 150 nm for HEK293FT cells, outperforming larger particles (245 nm) in terms of transfection efficiency. Similarly, McMillan et al. [21] investigated LNP-mediated mRNA expression in HEK293 cells and found an optimal size range of 105–138 nm for maximizing potency, which closely aligns with our simulation results.

**Fig. 8:**
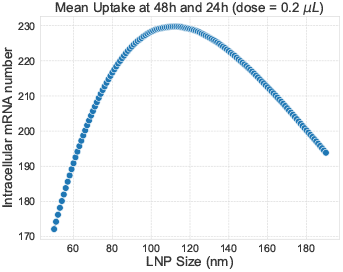
Mean intracellular mRNA uptake at 24 h and 48 h as a function of LNP size (dose = 0.2 *µL*).

### Influence of Cell Area on LNP Uptake Efficiency

The validated multi-scale kinetic model was used to conduct *in-sillico* experiments examining how cell area influences mRNA–LNP uptake. As illustrated in Figure 3D, both the magnitude of LNP uptake and its cell-to-cell variability show a positive correlation with cell area. This trend is consistent with prior observations in lipid-based and other nanoparticle delivery systems [76, 7, 8]. To investigate the mechanistic underpinnings of this correlation, we performed simulations and analyzed mRNA potency predictions across four quartiles of cell area (Q1–Q4), corresponding to the 0–25th, 25–50th, 50–75th, and 75–100th percentiles of the measured cell area distribution. The results are presented in Figure 7.

Simulation predictions of intracellular mRNA copy numbers closely match experimental observations from the smFISH assay for the first three cell area quartiles (Q1–Q3), with a slight overestimation observed in the largest quartile (Q4). These results suggest a positive correlation between cell area and mRNA uptake, though the relationship is sublinear. Notably, the developed multi-scale kinetic model captures this quantitative trend effectively. While larger cells may possess more surface receptors, enabling higher LNP uptake rates [8, 7], this advantage may be transient due to cell cycle dynamics. According to the cell area–dependent division model described in Eqs. (4)–(8), larger cells are more likely to enter mitosis, during which intracellular mRNA levels are diluted through partitioning between daughter cells.

The proposed framework enhances our understanding of how the internalization rate responds to changes in ligand concentration and cell area, ultimately approaching a saturation point. Monod kinetics is applied to model ligand diffusion and receptor-mediated internalization in Eq. (11), particularly in scenarios where LNP ligands bind to a finite number of cell surface receptors—a quantity that scales with cell area. Under this model, the empirical constant *K*_*A*_ modulates the sensitivity of the uptake rate to cell area. In addition, *K*_bind_ reflects the ligand–receptor binding affinity, thereby linking molecular interaction strength to macroscopic uptake behavior [77].

## Discussion

While mRNA vaccines—particularly those developed for COVID-19—have demonstrated high efficacy, challenges remain in optimizing their delivery and expression efficiency, especially in proliferating cells. Enhancing critical quality attributes (CQAs), such as lipid nanoparticle (LNP) size, is essential for maximizing the potency and therapeutic effectiveness of LNP-based mRNA delivery systems [78, 31, 79]. However, this optimization is often constrained by the high cost and time requirements associated with conventional experimental approaches. Furthermore, the influence of additional variables—such as cell density, proliferation rate, and dynamic changes in cell morphology—on mRNA-LNP potency remains insufficiently characterized and poorly understood.

The proposed MIMS addresses these challenges by capturing the underlying mechanisms of the mRNA-LNP delivery and expression process. This model accounts for the interactive effects of multiple factors on potency across molecular, cellular, and macroscopic scales. By integrating with advanced assays (such as smFISH and flow cytometry) capable of quantifying gene and protein expression at the single-cell level, MIMS enhances mechanistic understanding and enables robust potency prediction. Importantly, it provides a predictive distribution that captures cell-to-cell variability, supporting the development of rapid and reliable quality control strategies for mRNA-LNP formulations.

This multi-scale kinetic model offers mechanistic insights into how CQAs influence mRNA-LNP potency and contribute to cell-to-cell variability. While prior studies have investigated variability in delivery efficiency, they have not fully elucidated the underlying sources of heterogeneity observed in *in vitro* potency assays. For instance, Leonhardt *et al*. [12] proposed a statistical framework for modeling mRNA-lipoplex delivery, attributing cell-to-cell variability in uptake primarily to stochastic fluctuations in endosomal loading and lysis. Similarly, Rees *et al*. [7] developed a probabilistic model to examine the effects of cell membrane properties and dosage on delivery variability. However, their approaches lack mechanistic interpretability of the LNP binding process, leaving key stochastic drivers of this variability uncharacterized.

The validated MIMS model identified LNP size as a key contributor to cell-to-cell variability in uptake, particularly through its effects on internalization kinetics and binding efficiency. Both theoretical [5, 6, 16] and experimental studies [80, 81] have shown that internalization time exhibits a V-shaped relationship with LNP size: as size increases beyond a lower threshold, internalization time decreases until reaching an optimal size, after which it increases again. While this size-dependent internalization behavior is well established at the single-particle level, *in vitro* LNP uptake is also modulated by dose and cell density dynamics. Additionally, intracellular mRNA copy number is influenced by the mRNA loading per LNP [82]. The MIMS model captures the combined effects of these factors on mRNA-LNP uptake. Larger LNPs can encapsulate more mRNA [18, 19]; however, at a fixed mRNA dose, this results in fewer total particles, reducing the number of LNP-receptor binding events. This mechanistic insight explains the reduced transfection efficiency observed by Liao et al. [19] for high-loading LNPs at constant mRNA dose, as well as the lower long-term antibody titers reported in mice administered larger LNPs [75].

The developed MIMS model advances quantitative understanding of how various sources of uncertainty contribute to cell-to-cell variability in potency during *in vitro* mRNA-LNP delivery. Intracellular mRNA copy number increases progressively with LNP uptake, but is simultaneously reduced by three key processes: (1) dilution due to cell division, (2) exocytosis of mRNA, and (3) mRNA degradation. The stochastic nature of each of these processes further amplifies variability in potency. While uptake and degradation have been quantitatively modeled in prior studies [12, 39, 40], the contributions of cell division and exocytosis remain largely unquantified. Commonly used cell lines in mRNA-LNP studies—such as HEK293 and its derivatives [19, 20, 21, 22], as well as HeLa cells [23, 24, 25]—exhibit doubling times ranging from 24 to 45 hours [26, 27]. Therefore, for *in vitro* potency assessments extending beyond 24 hours, dilution of exogenous mRNA and its translated proteins becomes a significant contributor to variability. MIMS incorporates this effect through a cell division module, where cell area dynamics influence both receptor abundance and division rate, thereby modulating intracellular mRNA levels over time. This mechanistic insight also helps elucidate the role of the cell cycle in mRNA-LNP potency [7, 83, 84]. Moreover, as described in Eq. (13), proliferating cells continuously deplete extracellular mRNA-LNPs through uptake, introducing a time-dependent reduction in available LNPs—a factor rarely considered in previous models.

This study opens several promising avenues for future research. One key direction is the quantitative characterization of ligand–receptor binding affinity, which governs the empirical coefficient *K*_bind_ in the Monod model describing mRNA-LNP uptake kinetics. Binding affinity is influenced by multiple interacting factors, including ligand and receptor densities, LNP surface charge, and lipid composition. A systematic investigation into how these physicochemical parameters and CQAs affect the dynamics and variability of the uptake process could deepen mechanistic understanding and inform the optimal design and quality control of mRNA-LNP products. Another important area is the regulation of protein expression—a central determinant of potency—through improved characterization of translation rates and mRNA degradation kinetics across different mRNA sequences. Understanding how mRNA structural features influence RNA stability and protein synthesis would significantly enhance the mechanistic modeling of post-delivery gene expression. Additionally, the role of cell lines in modulating mRNA-LNP potency warrants further exploration. Prior studies have shown that identical mRNA-LNP formulations can elicit markedly different responses across cell types [24, 21]. While our model accounts for cell density and area dynamics, other factors—such as membrane elasticity, receptor specificity, translational capacity, and mRNA degradation rates—may also contribute to cell line–dependent variability. Future work should aim to quantify these contributions to better support the development of cell-targeted mRNA-LNP delivery systems.

## Data Availability

All the experiment data and code in this paper can be accessed once accepted.

## Supplementary Data statement

Supplementary Data are available online.

## Author contributions

Yuling Yang (Conceptualization [lead], Formal analysis [lead], Investigation [equal], Methodology [lead], Software [equal], Visualization [lead], Writing - original draft [lead]), Yuchen Qiu (Data curation [lead], Investigation [equal], Software [equal], Validation [lead], Visualization [lead], Writing - original draft [equal]), Keqi Wang (Formal analysis [equal], Methodology [lead], Writing - review & editing [equal]), Yifang Liu (Data curation [equal], Investigation [equal], Software [equal], Validation [equal]), Gautam Sanyal (Conceptualization [equal], Funding acquisition [lead]), Paul C. Whitford (Funding acquisition [equal], Writing - review & editing [equal]), Sara Rouhanifard (Conceptualization [lead], Data curation [equal], Funding acquisition [equal], Project administration [equal], Resources [lead], Supervision [lead], Writing - review & editing [equal]), Wei Xie (Conceptualization [lead], Methodology [equal], Project administration [lead], Resources [equal], Supervision [lead], Writing - review & editing [lead]).

## References

1. M. D. Buschmann, M. J. Carrasco, S. Alishetty, M. Paige, M. G. Alameh, and D. Weissman. Nanomaterial Delivery Systems for mRNA Vaccines. Vaccines, 9(1), January 2021.

2. Olga Vasileva, Olga Zaborova, Bogdan Shmykov, Roman Ivanov, and Vasiliy Reshetnikov. Composition of lipid nanoparticles for targeted delivery: application to mrna therapeutics. Frontiers in Pharmacology, 15:1466337, 2024.

3. G. Sanyal. Development of functionally relevant potency assays for monovalent and multivalent vaccines delivered by evolving technologies. Npj Vaccines, 7(1), May 2022.

4. Valentina Francia, Keni Yang, Sarah Deville, Catharina Reker-Smit, Inge Nelissen, and Anna Salvati. Corona Composition Can Affect the Mechanisms Cells Use to Internalize Nanoparticles. ACS Nano, 13(10):11107–11121, October 2019.

5. H. Gao, W. Shi, and L. B. Freund. Mechanics of receptor-mediated endocytosis. Proc Natl Acad Sci U S A, 102(27):9469–74, July 2005.

6. S. Zhang, H. Gao, and G. Bao. Physical Principles of Nanoparticle Cellular Endocytosis. ACS Nano, 9(9):8655–71, September 2015.

7. P. Rees, J. W. Wills, M. R. Brown, C. M. Barnes, and H. D. Summers. The origin of heterogeneous nanoparticle uptake by cells. Nature Communications, 10(1):2341, May 2019.

8. C. Aberg, V. Piattelli, D. Montizaan, and A. Salvati. Sources of variability in nanoparticle uptake by cells. Nanoscale, 13(41):17530–17546, October 2021.

9. Jonathan L. Kirschman, Sushma Bhosle, Daryll Vanover, Emmeline L. Blanchard, Kristin H. Loomis, Chiara Zurla, Kathryn Murray, Blaine C. Lam, and Philip J. Santangelo. Characterizing exogenous mRNA delivery, trafficking, cytoplasmic release and RNA–protein correlations at the level of single cells. Nucleic acids research, 45(12):e113–e113, 2017.

10. Cory D. Sago, Melissa P. Lokugamage, Kalina Paunovska, Daryll A. Vanover, Christopher M. Monaco, Nirav N. Shah, Marielena Gamboa Castro, Shannon E. Anderson, Tobi G. Rudoltz, Gwyneth N. Lando, Pooja Munnilal Tiwari, Jonathan L. Kirschman, Nick Willett, Young C. Jang, Philip J. Santangelo, Anton V. Bryksin, and James E. Dahlman. High-throughput in vivo screen of functional mRNA delivery identifies nanoparticles for endothelial cell gene editing. Proceedings of the National Academy of Sciences, 115(42):E9944–E9952, October 2018.

11. Kangzeng Wu, Fengwei Xu, Yongchao Dai, Shanshan Jin, Anjie Zheng, Ning Zhang, and Yuhong Xu. Characterization of mrna-lnp structural features and mechanisms for enhanced mrna vaccine immunogenicity. Journal of Controlled Release, 376:1288–1299, 2024.

12. C. Leonhardt, G. Schwake, T. R. Stogbauer, S. Rappl, J. T. Kuhr, T. S. Ligon, and J. O. Radler. Single-cell mRNA transfection studies: Delivery, kinetics and statistics by numbers. Nanomedicine, 10(4):679–88, May 2014.

13. H. Yasar, A. Biehl, C. De Rossi, M. Koch, X. Murgia, B. Loretz, and C. M. Lehr. Kinetics of mRNA delivery and protein translation in dendritic cells using lipid-coated PLGA nanoparticles. J Nanobiotechnology, 16(1):72, September 2018.

14. Andre E. Nel, Lutz Mädler, Darrell Velegol, Tian Xia, Eric M. V. Hoek, Ponisseril Somasundaran, Fred Klaessig, Vince Castranova, and Mike Thompson. Understanding biophysicochemical interactions at the nano–bio interface. Nature Materials, 8(7):543–557, July 2009.

15. Gang Bao and X Robert Bao. Shedding light on the dynamics of endocytosis and viral budding. Proceedings of the National Academy of Sciences, 102(29):9997–9998, 2005.

16. Hongyan Yuan and Sulin Zhang. Effects of particle size and ligand density on the kinetics of receptor-mediated endocytosis of nanoparticles. Applied Physics Letters, 96(3), 2010.

17. Thomas S. Ligon, Carolin Leonhardt, and Joachim O. Rädler. Multi-Level Kinetic Model of mRNA Delivery via Transfection of Lipoplexes. PLOS ONE, 9(9):e107148, September 2014.

18. V. P. Zhdanov. Kinetics of lipid-nanoparticle-mediated intracellular mRNA delivery and function. Physical Review E, 96(4), October 2017.

19. S. Liao, S. Wang, A. Wadhwa, A. Birkenshaw, K. Fox, M. H. Y. Cheng, I. C. Casmil, A. A. Magana, N. V. Bathula, C. H. Ho, J. Y. Cheng, L. J. Foster, K. W. Harder, C. J. D. Ross, P. R. Cullis, and A. K. Blakney. Transfection Potency of Lipid Nanoparticles Containing mRNA Depends on Relative Loading Levels. ACS Appl Mater Interfaces, 17(2):3097–3105, January 2025.

20. J. Long, C. Yu, H. Zhang, Y. Cao, Y. Sang, H. Lu, Z. Zhang, X. Wang, H. Wang, G. Song, J. Yang, and S. Wang. Novel Ionizable Lipid Nanoparticles for SARS-CoV-2 Omicron mRNA Delivery. Adv Healthc Mater, 12(13):e2202590, May 2023.

21. Caitlin McMillan, Amy Druschitz, Stephen Rumbelow, Ankita Borah, Burcu Binici, Zahra Rattray, and Yvonne Perrie. Tailoring lipid nanoparticle dimensions through manufacturing processes. Rsc Pharmaceutics, 1(4):841–853, 2024.

22. J. Walther, D. Porenta, D. Wilbie, C. Seinen, N. Benne, Q. Yang, O. G. de Jong, Z. Lei, and E. Mastrobattista. Comparative analysis of lipid Nanoparticle-Mediated delivery of CRISPR-Cas9 RNP versus mRNA/sgRNA for gene editing in vitro and in vivo. Eur J Pharm Biopharm, 196:114207, March 2024.

23. N. Patel, Z. Davis, C. Hofmann, J. Vlasak, J. W. Loughney, P. DePhillips, and M. Mukherjee. Development and Characterization of an In Vitro Cell-Based Assay to Predict Potency of mRNA-LNP-Based Vaccines. Vaccines, 11(7), July 2023.

24. S. Patel, N. Ashwanikumar, E. Robinson, Y. Xia, C. Mihai, J. P. Griffith, S. Hou, A. A. Esposito, T. Ketova, K. Welsher, J. L. Joyal, O. Almarsson, and G. Sahay. Naturally-occurring cholesterol analogues in lipid nanoparticles induce polymorphic shape and enhance intracellular delivery of mRNA. Nature Communications, 11(1):983, February 2020.

25. M. J. Munson, G. O’Driscoll, A. M. Silva, E. Lazaro-Ibanez, Gallud, J. T. Wilson, A. Collen, E. K. Esbjorner, and A. Sabirsh. A high-throughput Galectin-9 imaging assay for quantifying nanoparticle uptake, endosomal escape and functional RNA delivery. Commun Biol, 4(1):211, February 2021.

26. Philip Thomas and Trevor G Smart. Hek293 cell line: a vehicle for the expression of recombinant proteins. Journal of pharmacological and toxicological methods, 51(3):187–200, 2005.

27. Lei Tang. Investigating heterogeneity in hela cells. Nature Methods, 16(4):281–281, 2019.

28. Dann Huh and Johan Paulsson. Random partitioning of molecules at cell division. Proceedings of the National Academy of Sciences, 108(36):15004–15009, 2011.

29. Namit Chaudhary, Drew Weissman, and Kathryn A. Whitehead. mRNA vaccines for infectious diseases: Principles, delivery and clinical translation. Nature Reviews Drug Discovery, 20(11):817–838, November 2021.

30. M. Maugeri, M. Nawaz, A. Papadimitriou, A. Angerfors, Camponeschi, M. Na, M. Hölttä, P. Skantze, S. Johansson, M. Sundqvist, J. Lindquist, T. Kjellman, I. L. Mårensson, T. Jin, P. Sunnerhagen, S. Östman\, L. Lindfors, and H. Valadi. Linkage between endosomal escape of LNP-mRNA and loading into EVs for transport to other cells. Nature Communications, 10, September 2019.

31. S. Douka, L. E. Brandenburg, C. Casadidio, J. Walther, B. B. M. Garcia, J. Spanholtz, M. Raimo, W. E. Hennink, E. Mastrobattista, and M. Caiazzo. Lipid nanoparticle-mediated messenger RNA delivery for ex vivo engineering of natural killer cells. J Control Release, 361:455–469, September 2023.

32. Changrong Wang, Caiyan Zhao, Weipeng Wang, Xiaoqing Liu, and Hongzhang Deng. Biomimetic noncationic lipid nanoparticles for mrna delivery. Proceedings of the National Academy of Sciences, 120(51):e2311276120, 2023.

33. Chuanmei Tang, Yexi Zhang, Bowen Li, Xiangwei Fan, Zixuan Wang, Rongxin Su, Wei Qi, and Yuefei Wang. Modular design of lipopeptide-based organ-specific targeting (post) lipid nanoparticles for highly efficient rna delivery. Advanced Materials, 37(11):2415643, 2025.

34. Jia Huang, Xin Bai, William Stewart, Xiaoyang Xu, and Xue-Qing Zhang. Budesonide-incorporated inhalable lipid nanoparticles for antitslp nanobody mrna delivery to treat steroid-resistant asthma. Nature Communications, 16(1):6013, 2025.

35. Alexandra Murschhauser, Peter J. F. Röttgermann, Daniel Woschée, Martina F. Ober, Yan Yan, Kenneth A. Dawson, and Joachim O. Rädler. A high-throughput microscopy method for single-cell analysis of event-time correlations in nanoparticle-induced cell death. Communications Biology, 2(1):35, January 2019.

36. Rafa-l Krzysztoń, Daniel Woschée, Anita Reiser, Gerlinde Schwake, Helmut H. Strey, and Joachim O. Rädler. Single-cell kinetics of siRNA-mediated mRNA degradation. Nanomedicine: Nanotechnology, Biology and Medicine, 21:102077, October 2019. remark: live-cell assay.

37. Kempe Simon Maximilian Reiser Anita, Woschée Daniel. Live-cell imaging of single-cell arrays (LISCA) - a versatile technique to quantify cellular kinetics. JoVE, (169):e62025, 2021.

38. H. Parhiz, V. V. Shuvaev, Q. Li, T. E. Papp, A. A. Akyianu, R. Shi, A. Yadegari, H. Shahnawaz, S. C. Semple, B. L. Mui, D. Weissman, V. R. Muzykantov, and P. M. Glassman. Physiologically based modeling of LNP-mediated delivery of mRNA in the vascular system. Mol Ther Nucleic Acids, 35(2):102175, June 2024.

39. R. Mihaila, D. Ruhela, E. Keough, E. Cherkaev, S. Chang, B. Galinski, R. Bartz, D. Brown, B. Howell, and J. J. Cunningham. Mathematical Modeling: A Tool for Optimization of Lipid Nanoparticle-Mediated Delivery of siRNA. Mol Ther Nucleic Acids, 7:246–255, June 2017.

40. R. Mihaila, D. Ruhela, B. Galinski, A. Card, M. Cancilla, T. Shadel, J. Kang, S. Tep, J. Wei, R. M. Haas, J. Caldwell, W. M. Flanagan, N. Kuklin, E. Cherkaev, and B. Ason. Modeling the Kinetics of Lipid-Nanoparticle-Mediated Delivery of Multiple siRNAs to Evaluate the Effect on Competition for Ago2. Mol Ther Nucleic Acids, 16:367–377, June 2019.

41. Huw D. Summers, Paul Rees, Mark D. Holton, M. Rowan Brown, Sally C. Chappell, Paul J. Smith, and Rachel J. Errington. Statistical analysis of nanoparticle dosing in a dynamic cellular system. Nature Nanotechnology, 6(3):170–174, March 2011.

42. Geoffrey M. Cooper and Kenneth Adams. The Cell: A Molecular Approach. Oxford University Press, 2022.

43. Katharina Schiessl, Swathi Kausika, Paul Southam, Max Bush, and Robert Sablowski. Jagged controls growth anisotropy and coordination between cell size and cell cycle during plant organogenesis. Current Biology, 22(19):1739–1746, 2012.

44. Helmut Dolznig, Florian Grebien, Thomas Sauer, Hartmut Beug, and Ernst W Müllner. Evidence for a size-sensing mechanism in animal cells. Nature cell biology, 6(9):899–905, 2004.

45. D Killander and A Zetterberg. Quantitative cytochemical studies on interphase growth: I. determination of dna, rna and mass content of age determined mouse fibroblasts in vitro and of intercellular variation in generation time. Experimental cell research, 38(2):272–284, 1965.

46. W. J. Thieman and M. A. Palladino. Introduction to Biotechnology. Pearson Education, 2012.

47. Conor M O’Brien, Qi Zhang, Prodromos Daoutidis, and Wei-Shou Hu. A hybrid mechanistic-empirical model for in silico mammalian cell bioprocess simulation. Metabolic Engineering, 66:31–40, 2021.

48. Keqi Wang, Sarah W Harcum, and Wei Xie. Multi-scale kinetics modeling for cell culture process with metabolic state transition. arXiv preprint arXiv:2412.03883, 2024.

49. Eleni Mantikou, Kai Mee Wong, Sjoerd Repping, and Sebastiaan Mastenbroek. Molecular origin of mitotic aneuploidies in preimplantation embryos. Biochimica et Biophysica Acta (BBA)-Molecular Basis of Disease, 1822(12):1921–1930, 2012.

50. G. I. Bell and E. C. Anderson. Cell growth and division. I. A mathematical model with applications to cell volume distributions in mammalian suspension cultures. Biophys J, 7(4):329–51, July 1967.

51. R. Jones A, M. Forero-Vargas, S. P. Withers, R. S. Smith, J. Traas, W. Dewitte, and J. A. H. Murray. Cell-size dependent progression of the cell cycle creates homeostasis and flexibility of plant cell size. Nature Communications, 8:15060, April 2017.

52. Chen Jia, Abhyudai Singh, and Ramon Grima. Concentration fluctuations in growing and dividing cells: Insights into the emergence of concentration homeostasis (with cell area increase, the protein synthesis rate should increase to maintain constant concentration). PLoS computational biology, 18(10):e1010574, 2022.

53. Amit Tzur, Ran Kafri, Valerie S LeBleu, Galit Lahav, and Marc W Kirschner. Cell growth and size homeostasis in proliferating animal cells. Science, 325(5937):167–171, 2009.

54. A. Provost, G. Bastin, S. N. Agathos, and Y. J. Schneider. Metabolic design of macroscopic bioreaction models: Application to Chinese hamster ovary cells. Bioprocess Biosyst Eng, 29(5-6):349–66, December 2006.

55. S. Tzlil, M. Deserno, W. M. Gelbart, and A. Ben-Shaul. A statistical-thermodynamic model of viral budding. Biophys J, 86(4):2037–48, April 2004.

56. W. Rawicz, K. C. Olbrich, T. McIntosh, D. Needham, and E. Evans. Effect of Chain Length and Unsaturation on Elasticity of Lipid Bilayers. Biophysical Journal, 79(1):328–339, July 2000.

57. S. Ashraf, A. H. Said, R. Hartmann, M. A. Assmann, N. Feliu, P. Lenz, and W. J. Parak. Quantitative Particle Uptake by Cells as Analyzed by Different Methods. Angewandte Chemie-International Edition, 59(14):5438–5453, March 2020.

58. Jawahar Khetan, Md Shahinuzzaman, Sutapa Barua, and Dipak Barua. Quantitative analysis of the correlation between cell size and cellular uptake of particles. Biophysical Journal, 116(2):347–359, 2019.

59. Tania López-Herńandez, Volker Haucke, and Tanja Maritzen. Endocytosis in the adaptation to cellular stress. Cell stress, 4(10):230, 2020.

60. Costin N Antonescu, Timothy E McGraw, and Amira Klip. Reciprocal regulation of endocytosis and metabolism. Cold Spring Harbor perspectives in biology, 6(7):a016964, 2014.

61. S. S. Vallender. Calculation of the wasserstein distance between probability distributions on the line. Theory of Probability & Its Applications, 18(4):784–786, 1974.

62. Diederik P Kingma and Jimmy Ba. Adam: A method for stochastic optimization. arXiv preprint arXiv:1412.6980, 2014.

63. John Wilder Tukey et al. Exploratory data analysis, volume 2. Springer, 1977.

64. M. Monici. Cell and tissue autofluorescence research and diagnostic applications. Biotechnol Annu Rev, 11:227–56, 2005.

65. Zhi Xiong Chong, Swee Keong Yeap, and Wan Yong Ho. Transfection types, methods and strategies: a technical review. PeerJ, 9:e11165, 2021.

66. Giuditta Guerrini, Diletta Scaccabarozzi, Dora Mehn, Ambra Sarracino, Sabrina Gioria, and Luigi Calzolai. Challenges in measuring in vitro activity of lnp-mrna therapeutics. International Journal of Molecular Sciences, 26(17):8152, 2025.

67. Sarah Lindsay, Muattaz Hussain, Burcu Binici, and Yvonne Perrie. Exploring the challenges of lipid nanoparticle development: the in vitro–in vivo correlation gap. Vaccines, 13(4):339, 2025.

68. Hong Sun, Songyu Wang, Mei Lu, Christine E Tinberg, and Benjamin M Alba. Protein production from hek293 cell line-derived stable pools with high protein quality and quantity to support discovery research. PLoS One, 18(6):e0285971, 2023.

69. Audrey Le Ru, Danielle Jacob, Julia Transfiguracion, Sven Ansorge, Olivier Henry, and Amine A Kamen. Scalable production of influenza virus in hek-293 cells for efficient vaccine manufacturing. Vaccine, 28(21):3661–3671, 2010.

70. Emma Petiot, Danielle Jacob, Stephane Lanthier, Verena Lohr, Sven Ansorge, and Amine A Kamen. Metabolic and kinetic analyses of influenza production in perfusion hek293 cell culture. BMC biotechnology, 11:1–12, 2011.

71. E Trucco. On the average cellular volume in synchronized cell populations. The bulletin of mathematical biophysics, 32:459–473, 1970.

72. Peter A Fantes, WD Grant, RH Pritchard, PE Sudbery, and AE Wheals. The regulation of cell size and the control of mitosis. Journal of theoretical biology, 50(1):213–244, 1975.

73. X. Jiang, C. Rocker, M. Hafner, S. Brandholt, R. M. Dorlich, and G. U. Nienhaus. Endo- and exocytosis of zwitterionic quantum dot nanoparticles by live HeLa cells. ACS Nano, 4(11):6787–97, November 2010.

74. H. Shen, G. Nelson, D. E. Nelson, S. Kennedy, D. G. Spiller, T. Griffiths, N. Paton, S. G. Oliver, M. R. White, and D. B. Kell. Automated tracking of gene expression in individual cells and cell compartments. J R Soc Interface, 3(11):787–94, December 2006.

75. K. J. Hassett, J. Higgins, A. Woods, B. Levy, Y. Xia, C. J. Hsiao, E. Acosta, O. Almarsson, M. J. Moore, and L. A. Brito. Impact of lipid nanoparticle size on mRNA vaccine immunogenicity. J Control Release, 335:237–246, July 2021.

76. Xinlong Wang, Xiaohong Hu, Jingchao Li, Adriana C Mulero Russe, Naoki Kawazoe, Yingnan Yang, and Guoping Chen. Influence of cell size on cellular uptake of gold nanoparticles. Biomaterials science, 4(6):970–978, 2016.

77. Mostafa Maalmi, William Strieder, and Arvind Varma. Ligand diffusion and receptor mediated internalization: Michaelis– menten kinetics. Chemical engineering science, 56(19):5609–5616, 2001.

78. Burcu Binici, Zahra Rattray, Assaf Zinger, and Yvonne Perrie. Exploring the impact of commonly used ionizable and pegylated lipids on mRNA-LNPs: A combined in vitro and preclinical perspective. Journal of Controlled Release, 377:162–173, January 2025.

79. Marianna Yanez Arteta, Tomas Kjellman, Stefano Bartesaghi, Simonetta Wallin, Xiaoqiu Wu, Alexander J Kvist, Aleksandra Dabkowska, Noémi Székely, Aurel Radulescu, Johan Bergenholtz, et al. Successful reprogramming of cellular protein production through mrna delivered by functionalized lipid nanoparticles. Proceedings of the National Academy of Sciences, 115(15):E3351–E3360, 2018.

80. Yasuhiro Aoyama, Takuya Kanamori, Takashi Nakai, Toshinori Sasaki, Shohei Horiuchi, Shinsuke Sando, and Takuro Niidome. Artificial viruses and their application to gene delivery. Size-controlled gene coating with glycocluster nanoparticles. Journal of the American Chemical Society, 125(12):3455–3457, 2003.

81. F. Osaki, T. Kanamori, S. Sando, T. Sera, and Y. Aoyama. A quantum dot conjugated sugar ball and its cellular uptake. On the size effects of endocytosis in the subviral region. J Am Chem Soc, 126(21):6520–1, June 2004.

82. S. Li, Y. Hu, A. Li, J. Lin, K. Hsieh, Z. Schneiderman, P. Zhang, Y. Zhu, C. Qiu, E. Kokkoli, T. H. Wang, and H. Q. Mao. Payload distribution and capacity of mRNA lipid nanoparticles. Nature Communications, 13(1):5561, September 2022.

83. J. A. Kim, C. Aberg, A. Salvati, and K. A. Dawson. Role of cell cycle on the cellular uptake and dilution of nanoparticles in a cell population. Nat Nanotechnol, 7(1):62–8, November 2011.

84. Einat Panet, Tal Mashriki, Roxane Lahmi, Abraham Jacob, Efrat Ozer, Manuela Vecsler, Itay Lazar, and Amit Tzur. The interface of nanoparticles with proliferating mammalian cells. Nature Nanotechnology, 12(7):598–600, 2017.

