## supplementaryMaterials for "Multi-Scale Kinetics Modeling and Advanced Assay for mRNA-Lipid Nanoparticle Potency Assessment"

### Supplementary Data

### 1 Derivation of Cell Area–Dependent Division Model

#### 1.1 Cell Density Growth Rate and Cell Cycle Transiton Rate

In this section, we derive the relationship between the transition rate $r_{21}(t_h)$, which governs the progression from the G1-S phase to the G2-M phase, and the cell density growth rate $\mu(t_h)$. At any time $t_h$, let $I^{div}(t_h) \in \{0 : \text{Not divide}, 1 : \text{divide}, -1 : \text{die}\}$ denote the indicator variable representing the division status of a single cell within a short time interval $\Delta t$. Let $z(t_h)$ denote the cell cycle state: if the cell is in G1-S phase, we have $z(t_h) \in$ G1-S, and $z(t_h) \in$ G2-M otherwise. The dynamics of cell density can then be expressed as:

$$\Delta X(t_h) = \sum_{j=1}^{X(t_h)} I_j^{div}(t_h),$$

where $X(t_h)$ is the total number of cells at time $t_h$ and $I_j^{div}(t_h)$ indicates the division event of the $j$-th cell. In the proposed framework, the transition probability from the G2-M phase to the G1-S phase phase is modeled by an exponential distribution:

$$P_{21} = P(\text{G2-M} \rightarrow \text{G1-S} \mid z(t_h) \in \text{G2-M}) = 1 - e^{-r_{21}(t_h)\Delta t},$$

where $r_{21}(t_h)$ denotes the transition rate from the G2-M phase to the G1-S phase at time $t_h$. Since only cells in the G2-M phase are capable of division, the conditional expectation of the change in cell density becomes:

$$\begin{aligned}\mathbb{E}[\Delta X(t_h) \mid X(t_h)] = \mathbb{E}\left[\sum_{j=1}^{X(t_h)} I_j^{div}(t_h) \middle| X(t_h)\right] \\ = \quad & P\left(\text{G2-M} \rightarrow \text{G1-S} \mid z(t_h) \in \text{G2-M}\right) P\left(z(t_h) \in \text{G2-M}\right) X(t_h) \\ = \quad & (1 - e^{-r_{21}(t_h)\Delta t})\gamma_{\text{G2-M}} X(t_h).\end{aligned}$$

Taking the limit as $\Delta t \rightarrow 0$, we obtain:

$$\lim_{\Delta t \rightarrow 0} \frac{\mathbb{E}[\Delta X(t_h) \mid X(t_h)]}{\Delta t} = \lim_{\Delta t \rightarrow 0} \frac{(1 - e^{-r_{21}(t_h)\Delta t})}{\Delta t} \gamma_{\text{G2-M}} X(t_h) \approx r_{21}(t_h)\gamma_{\text{G2-M}} X(t_h).$$

Alternatively, the expected change in cell density can be expressed as: $\mathbb{E}\left[\Delta X(t_h) \mid X(t_h)\right] = \mu(t_h) X(t_h) \Delta t$, where $\mu(t_h)$ denotes the cell density growth rate at time $t_h$. By equating this expression with the previously derived expectation based on cell division dynamics and solving for $r_{21}$, we obtain:

$$r_{21}(t_h) \approx \frac{\mu(t_h)}{\gamma_{\text{G2-M}}}, \tag{S1}$$

where $\gamma_{\text{G2-M}}$ represents the proportion of living cells in the G2-M phase. This relationship links the transition rate from G2-M to G1-S with the overall cell density growth rate.

#### 1.2 Transition Probability from G1-S to G2-M Phase

We derive the expected transition probability from the G1-S phase to the G2-M phase, denoted by $P_{12}$. Let $\#$(G1-S)$(t_h)$ denote the number of cells in the G1-S phase at time $t_h$. Given the total cell density $X(t_h)$, the expected number of cells in the G1-S phase at time $t_h + \Delta t$ is:

$$
\begin{aligned}
&\mathbb{E}\left[\#(\text{G1-S})(t_h+\Delta t) \mid X(t_h)\right] \\
&= \mathbb{E}\Bigg[\sum_{i=1}^{X(t_h)} I(z_i(t_h) \in \text{G1-S})\left(I(z_i(t_h+\Delta t) \in \text{G1-S}) - I(z_i(t_h+\Delta t) \in \text{G2-M})\right) \\
&\quad +2\sum_{i=1}^{X(t_h)} I(z_i(t_h) \in \text{G2-M}) I(z_j(t_h+\Delta t) \in \text{G1-S})\Bigg] \qquad \text{(S2)} \\
&= \gamma_{\text{G1-S}} X(t_h)(1-2P_{12}) + 2\gamma_{\text{G2-M}} X(t_h)(1-e^{-r_{21}(t_h)\Delta t}).
\end{aligned}
$$

The second term in (S2) accounts for the doubling of cells due to mitosis at the end of the G2-M phase, followed by re-entry into the G1-S phase.

Similarly, the expected number of cells in the G2-M phase at time $t_h + \Delta t$ is:

$$
\begin{aligned}
&\mathbb{E}[\#(\text{G2-M})(t_h+\Delta t) \mid X(t_h)] \\
&= \mathbb{E}\Bigg[\sum_{i=1}^{X(t_h)} I(z_i(t_h) \in \text{G2-M})\left(I(z_i(t_h+\Delta t) \in \text{G2-M}) - I(z_i(t_h+\Delta t) \in \text{G1-S})\right) \\
&\quad +\sum_{i=1}^{X(t_h)} I(z_i(t_h) \in \text{G1-S}) I(z(t_h+\Delta t) \in \text{G2-M})\Bigg] \\
&= \gamma_{\text{G2-M}} X(t_h)(2e^{-r_{21}(t_h)\Delta t} - 1) + \gamma_{\text{G1-S}} X(t_h) P_{12}.
\end{aligned}
$$

Assuming the proportions of cells in each phase remain constant over time, we equate the ratio of expected cell counts:

$$
\frac{\gamma_{\text{G1-S}}}{\gamma_{\text{G2-M}}} = \frac{\gamma_{\text{G1-S}} X(t_h)(1-2P_{12}) + 2\gamma_{\text{G2-M}} X(t_h)(1-e^{-r_{21}(t_h)\Delta t})}{\gamma_{\text{G2-M}} X(t_h)(2e^{-r_{21}(t_h)\Delta t}-1) + \gamma_{\text{G1-S}} X(t_h) P_{12}}.
$$

Simplifying and solving for $P_{12}$, we obtain:

$$
\begin{aligned}
P_{12} &= P(\text{G1-S} \to \text{G2-M}) \\
&= \frac{2\gamma_{\text{G2-M}}(\gamma_{\text{G1-S}} + \gamma_{\text{G2-M}})}{\gamma_{\text{G1-S}}(\gamma_{\text{G1-S}} + 2\gamma_{\text{G2-M}})}(1-e^{-r_{21}(t_h)\Delta t}) \\
&= \frac{2\gamma_{\text{G2-M}}}{1-\gamma_{\text{G2-M}}^2}(1-e^{-r_{21}(t_h)\Delta t}).
\end{aligned}
$$

#### 1.3 Transition Probability from G1-S to G2-M Phase Conditioned on Cell Area

We derive the transition probability from the G1-S phase to the G2-M phase as a function of the cell area, denoted by $P_{12}(A(t_h) = a)$. Let $q(a)$ denote the stationary distribution density of cell area, and $g(t_h)$ denote the cell area growth rate at time $t_h$. Given a cell with area $A(t_h) = a$, the conditional transition probability from G1-S to G2-M over a short interval $\Delta t$, as described in Eq. (7) of the main text, is computed using Bayes' rule:

$$\begin{aligned}
&P(z(t_h + \Delta t) \in \text{G2-M} \mid z(t_h) \in \text{G1-S}, A(t_h) = a) \\
&= \frac{q(A(t_h) = a \mid z(t_h + \Delta t) \in \text{G2-M}, z(t_h) \in \text{G1-S}) P(z(t_h + \Delta t) \in \text{G2-M}, z(t_h) \in \text{G1-S})}{q(A(t_h) = a \mid z(t_h) \in \text{G1-S}) P(z(t_h) \in \text{G1-S})} \\
&= \frac{q(A(t_h) = a \mid z(t_h + \Delta t) \in \text{G2-M}, z(t_h) \in \text{G1-S})}{q(A(t_h) = a \mid z(t_h) \in \text{G1-S})} P_{12} \\
&= \frac{q(A(t_h + \Delta t) = a + g(t_h) a \Delta t \mid z(t_h + \Delta t) \in \text{G2-M})}{q(A(t_h) = a \mid z(t_h) \in \text{G1-S})} P_{12} \\
&= \frac{q(A(t_h + \Delta t) = a + g(t_h) a \Delta t \mid z(t_h + \Delta t) \in \text{G2-M})}{[q(A(t_h) = a) - q(A(t_h) = a, z(t_h) \in \text{G2-M})]/\gamma_{\text{G1-S}}} P_{12}. \\
&= \frac{(1 - \gamma_{\text{G2-M}})\, q(A(t_h + \Delta t) = a + g(t_h) a \Delta t \mid z(t_h + \Delta t) \in \text{G2-M})}{q(A(t_h) = a) - q(A(t_h) = a, z(t_h) \in \text{G2-M})} P_{12}. \\
&= \frac{(1 - \gamma_{\text{G2-M}})\, q(A(t_h + \Delta t) = a + g(t_h) a \Delta t \mid z(t_h + \Delta t) \in \text{G2-M})}{q(A(t_h) = a) - q(A(t_h) = a | z(t_h) \in \text{G2-M}) P(z(t_h) \in \text{G2-M})} P_{12}.
\end{aligned}$$

To evaluate the joint density $q(A(t_h) = a, z(t_h) \in \text{G2-M})$, we apply Bayes' rule:

$$q\left(a \mid z\left(t_h\right) \in \text{G2-M}\right) = \frac{q\left(z\left(t_h\right) \in \text{G2-M} | A\left(t_h\right) = a\right) q\left(a\right)}{\gamma_{\text{G2-M}}}.$$

Following the modeling approach in Provost et al. (2006), the conditional distribution $q\left(z\left(t_h\right) \in \text{G2-M} | A\left(t_h\right) = a\right)$ is represented by a sigmoid function:

$$P\left(z\left(t_h\right) \in \text{G2-M} | A\left(t_h\right) = a\right) = \text{Sigmoid}\left(\beta_1 a + \beta_0\right),$$

where $\beta_1 > 0$ and $\beta_0$ are the linear transformation coefficients of cell area. Substituting this into the joint density expression:

$$\begin{aligned}
q(A(t_h) = a, z(t_h) \in \text{G2-M}) &= P(z(t_h) \in \text{G2-M} | A(t_h) = a) q(a) \\
&= \text{Sigmoid}(\beta_1 a + \beta_0) q(a).
\end{aligned}$$

Finally, the area-dependent transition probability, as presented in Eq. (9) of the main text, is derived through the following steps:

$$\begin{aligned}
&P(z(t_h + \Delta t) \in \text{G2-M} \mid z(t_h) \in \text{G1-S}, A(t_h) = a) \\
&= \frac{(1 - \gamma_{\text{G2-M}})\, q(A(t_h + \Delta t) = a + g(t_h) a \Delta t \mid z(t_h + \Delta t) \in \text{G2-M})}{q(a) - \text{Sigmoid}(\beta_1 a + \beta_0) q(a)} P_{12} \\
&= \frac{(1 - \gamma_{\text{G2-M}})\, \text{Sigmoid}(\beta_1 (a + g(t_h) \Delta t) + \beta_0) q(a)}{(q(a) - \text{Sigmoid}(\beta_1 a + \beta_0) q(a)) \gamma_{\text{G2-M}}} P_{12} \\
&= \frac{(1 - \gamma_{\text{G2-M}})\, \text{Sigmoid}(\beta_1 (a + g(t_h) \Delta t) + \beta_0)}{(1 - \text{Sigmoid}(\beta_1 a + \beta_0)) \gamma_{\text{G2-M}}} P_{12}.
\end{aligned}$$

Table 1: Sequences of smFISH and qRT-PCR assays.

| Assay | Probe number | Sequence |
|---|---|---|
| smFISH | eGFP_1 | GTAAACAGTTCTTCGCCTTT |
| (5' - 3') | eGFP_2 | CATCCAGTTCCACCAGAATC |
| | eGFP_3 | ACGCTAAATTTATGGCCGTT |
| | eGFP_4 | TAAAGCACTGCACGCCATAG |
| | eGFP_5 | TTCATATGATCCGGATAGCG |
| | eGFP_6 | CTTCAAATTTCACTTCCGCG |
| | eGFP_7 | GGCTGTTATAGTTATATTCC |
| | eGFP_8 | ATCCGCCATAATATACACGT |
| | eGFP_9 | CTTTAATGCCGTTTTTCTGT |
| | eGFP_10 | CCATCTTCAATGTTATGGCG |
| | eGFP_11 | GGGTGTTCTGCTGATAATGA |
| | eGFP_12 | TCAGATAATGGTTATCCGGC |
| | eGFP_13 | TTTTTCGTTCGGATCTTTGC |
| | eGFP_14 | AAATTCCAGCAGCACCATAT |
| | eGFP_15 | TTATACAGTTCATCCATGCC |
| qRT-PCR | Forward | GCACAAGCTGGAGTACAACTA |
| (5' - 3') | Reverse | TGTTGTGGCGGATCTTGAA |
| | Probe | /56-FAM/AGCAGAAGA/ZEN/ACGGCATCAAGGTGA/3IABkFQ/ |

### 2 mRNA Sequences

Table 1 presents the sequences of smFISH and qRT-PCR assays. smFISH probes were designed using the Stellaris® Probe Designer (Biosearch Technologies). All probes were synthesized with a 3' end modification, mdC(TEG-Amino). Alexa Fluor™ 594 NHS Ester (Molecular Probes™, cat. no. A20004) was used for dye conjugation to the probes. qRT-PCR primers and probes were designed using the PrimerQuest™ Tool (Integrated DNA Technologies, IDT).

### 3 Cell Area Data Analysis

We analyzed the time-course cell area data, as shown in Figure S1. The distribution of cell areas was confirmed to follow a gamma distribution, demonstrated in Figures S1A and S1B. Figure S1C and Table 2 present the statistics of time-course cell area data. As illustrated in Figure S1D, the blue dots closely follow the line $y = x$. These pair-wise comparisons suggest that the the cell area distribution is time-invariant.

### References

Provost, A., Bastin, G., Agathos, S. N., and Schneider, Y. J. (2006). Metabolic design of macroscopic bioreaction models: Application to Chinese hamster ovary cells. *Bioprocess Biosyst Eng*, 29(5-6):349–66.

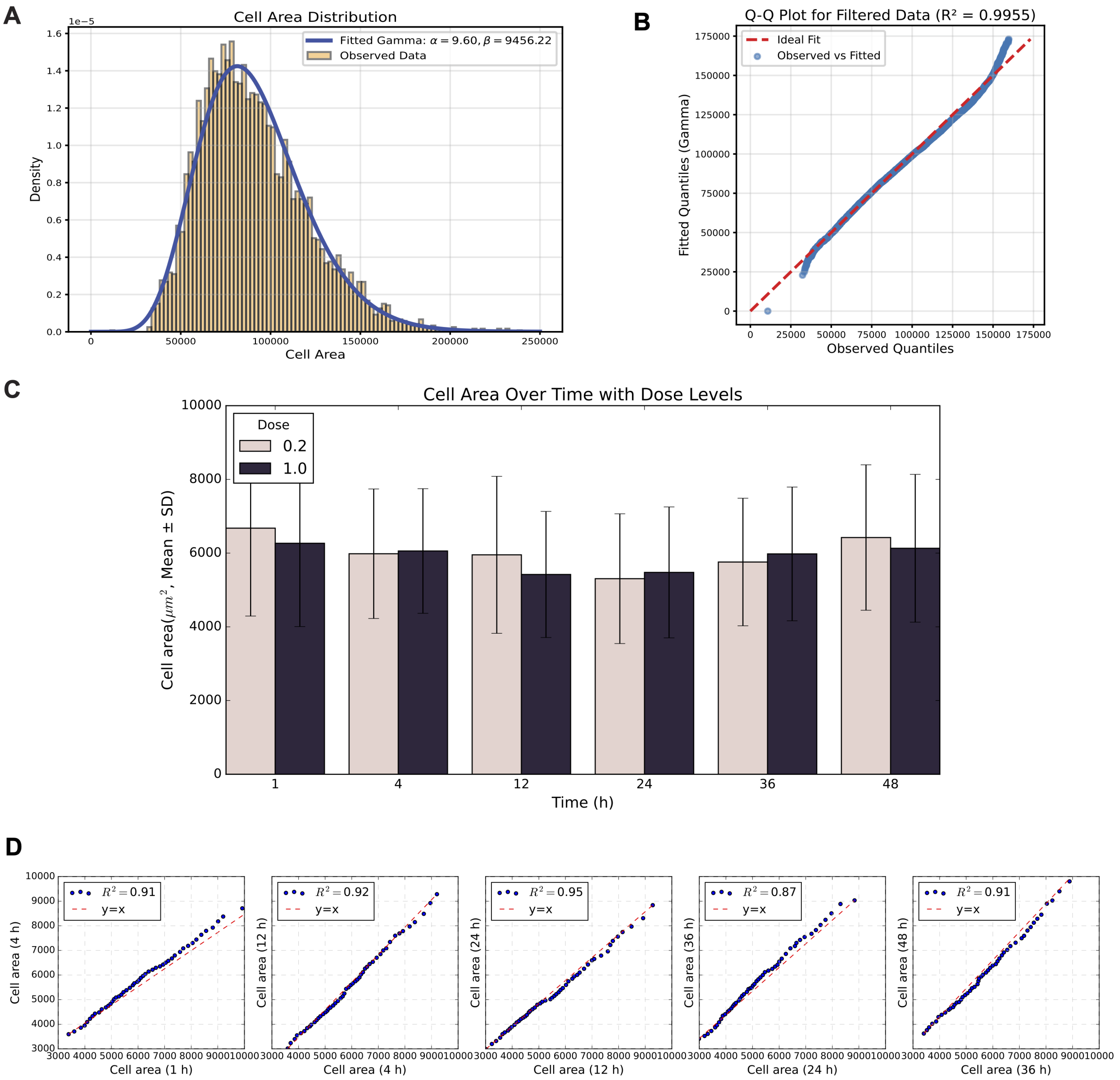


Figure S1: **Statistical tests of cell area.** (A) the distribution cell area; (B) Q-Q plot between observed cell area and fitted Gamma distribution. (C) Statistical test of time-course cell area; (D) Time-course Q-Q plots of observed cell area distributions.

Table 2: Statistics of cell area distribution.

| **Time (h)** | **Dose = 0.2** $\mu L$ | | **Dose = 1** $\mu L$ | |
|---|---|---|---|---|
| | Mean | Std | Mean | Std |
| 1 | 6677.33 | 2384.74 | 6267.53 | 2260.57 |
| 4 | 5983.16 | 1756.68 | 6057.98 | 1690.61 |
| 12 | 5953.92 | 2130.11 | 5421.06 | 1711.04 |
| 24 | 5306.72 | 1759.89 | 5477.13 | 1775.55 |
| 36 | 5758.69 | 1730.30 | 5979.10 | 1813.89 |
| 48 | 6422.89 | 1972.86 | 6131.75 | 2004.59 |
